# Convergent evolution of the SARS-CoV-2 Omicron subvariants leading to the emergence of BQ.1.1 variant

**DOI:** 10.1101/2022.12.05.519085

**Authors:** Jumpei Ito, Rigel Suzuki, Keiya Uriu, Yukari Itakura, Jiri Zahradnik, Sayaka Deguchi, Lei Wang, Spyros Lytras, Tomokazu Tamura, Izumi Kida, Hesham Nasser, Maya Shofa, MST Monira Begum, Masumi Tsuda, Yoshitaka Oda, Shigeru Fujita, Kumiko Yoshimatsu, Hayato Ito, Naganori Nao, Hiroyuki Asakura, Mami Nagashima, Kenji Sadamasu, Kazuhisa Yoshimura, Yuki Yamamoto, Tetsuharu Nagamoto, Gideon Schreiber, The Genotype to Phenotype Japan (G2P-Japan) Consortium, Akatsuki Saito, Keita Matsuno, Kazuo Takayama, Shinya Tanaka, Takasuke Fukuhara, Terumasa Ikeda, Kei Sato

## Abstract

In late 2022, although the SARS-CoV-2 Omicron subvariants have highly diversified, some lineages have convergently acquired amino acid substitutions at five critical residues in the spike protein. Here, we illuminated the evolutionary rules underlying the convergent evolution of Omicron subvariants and the properties of one of the latest lineages of concern, BQ.1.1. Our phylogenetic and epidemic dynamics analyses suggest that Omicron subvariants independently increased their viral fitness by acquiring the convergent substitutions. Particularly, BQ.1.1, which harbors all five convergent substitutions, shows the highest fitness among the viruses investigated. Neutralization assays show that BQ.1.1 is more resistant to breakthrough BA.2/5 infection sera than BA.5. The BQ.1.1 spike exhibits enhanced binding affinity to human ACE2 receptor and greater fusogenicity than the BA.5 spike. However, the pathogenicity of BQ.1.1 in hamsters is comparable to or even lower than that of BA.5. Our multiscale investigations provide insights into the evolutionary trajectory of Omicron subvariants.

## Introduction

As of November 2022, the SARS-CoV-2 Omicron variant (B. 1.1.529 and BA lineages) is the only current variant of concern (VOC)^1^. At the end of November 2021, Omicron BA.1 rapidly outcompeted Delta, a prior VOC. Soon after the global spread of Omicron BA.1 lineage, Omicron BA.2 became predominant in the world. Thereafter, a variety of BA.2 descendants, such as BA.5 and BA.2.75, emerged and are becoming predominant in certain countries.

Omicron BA.2 is highly diversified after emergence and is the origin of other recently emerging Omicron subvariants. Although both BA.5 and BA.2.75 diversified from BA.2, these two Omicron subvariants are phylogenetically independent to each other, suggesting that these two variants emerged independently^2^. However, recent studies including ours have demonstrated that the spike (S) proteins of these two variants exhibit similar evolutionary patterns: one is amino acid substitutions to evade antiviral humoral immunity, while the other is the substitution to increase the binding affinity to human angiotensin converting enzyme 2 (ACE2), the receptor for SARS-CoV-2 infection. For BA.5, the F486V substitution contributes to evasion from antiviral humoral immunity^3–6^, while the L452R substitution increases ACE2 binding affinity^4,7–9^. For BA.2.75, the G446S substitution is responsible for evasion from antiviral humoral immunity^6,10–14^, while the N460K substitution increases ACE2 binding affinity^2,7,11^. Because the substitutions in the S protein, that confer resistance to antiviral immunity, such as F486V and G446S, reduce the affinity to human ACE2^2,15^, it is conceivable to assume that the additional substitutions, such as L452R and N460K, compensate the decreased ACE2 binding affinity^2,15^.

Another common feature of BA.5 and BA.2.75 we found is the greater pathogenicity compared to BA.2 in a hamster model^2,15^. Because BA.5 and BA.2.75 are descendants of BA.2, our observations suggest that these two variants evolved to increase intrinsic pathogenicity. Importantly, our previous studies that focused on Delta^16^, Omicron BA.1^17^, Omicron BA.2^18^, Omicron BA.5^15^, and Omicron BA.2.75^2^ suggested that viral intrinsic pathogenicity in hamsters is closely associated with viral fusogenicity in cell culture system. Because previous studies suggest that higher fusogenicity is partly attributed to the increased affinity to human ACE2^11,15,18,19^, the substitutions in the S proteins of BA.5 (L452R) and BA.2.75 (N460K) can result in increasing intrinsic pathogenicity in hamsters.

Until the emergence of BA.5, the newly emerging SARS-CoV-2 variant globally outcompeted the previously predominant variant in a few months. However, as of November 2022, although a variety of Omicron BA.2 subvariants, including BA.2.75, have emerged after BA.5, none of them have successfully outcompeted BA.5 yet. Instead of the emergence of an outstanding SARS-CoV-2 variant, recently emerging Omicron subvariants are under convergent evolution: most variants have acquired substitutions at the same site of S, such as R346, K444, L452, N460, or F486. Notably, BQ. 1.1 is a descendant of BA.5 and bears all five recent convergent mutations: R346T, K444T, L452R, N460K, or F486V. As of October 12, 2022, the WHO classifies BQ.1.1 as an Omicron subvariant under monitoring^1^. Particularly, the BQ.1.1 S harbors two substitutions, L452R and N460K, that increase ACE2 binding affinity and fusogenicity^2,15^. These observations raise the possibility that BQ.1.1 is more fusogenic and pathogenic than BA.5. In this study, we illuminated the evolutionary principals underlying the current convergent evolution of Omicron lineages and characterized BQ.1.1 in terms of its transmissibility, immunogenicity, fusogenicity and intrinsic pathogenicity. Our results suggest that BQ.1.1 is a newly emerging variant that outcompete BA.5 and will be the globally predominant variant in the near future.

## Results

### Convergent evolution of Omicron lineages

As of November 2022, various Omicron lineages have continuously emerged, such as Omicron BA.1, BA.2, BA.4, BA.5, and BA.2.75 (**Fig. 1a**). As shown in **Fig. 1a**, BQ.1.1, a latest lineage of concern, emerged from the BA.5 cluster. Notably, the substitutions in S protein, particularly R346X, K444X, L452X, N460X, and F486X, seem to have convergently occurred in a variety of Omicron lineages (hereafter we refer to these five amino acid residues as “convergent sites” and substitutions at the residues as “convergent substitutions”)^6^. As described in the Introduction, the BQ.1.1 S harbors these five convergent substitutions, R346T, K444T, L452R, N460K, and F486V (**Fig. 1b, left**). Additionally, BQ.1.1 possesses six substitutions in the non-S region when compared to the parental BA.5 (**Fig. 1b, right**). To investigate the substitutions at these five convergent sites during Omicron evolution in depth, we constructed phylogenetic trees for BA.1, BA.2 (including BA.2.75), BA.4, and BA.5 (including BQ.1.1) and identified the branches on the trees where the convergent substitutions occurred (**Fig. 1c**). The R346 residue showed relatively higher substitution frequency compared to the other residues in all Omicron lineages (**Fig. 1c–e**). Consistent with our previous study^15^, the L452 residue in BA.2 showed the highest substitution frequency in that lineage (**Fig. 1c–e**). Importantly, the substitution events were more frequently detected in relatively younger lineages such as BA.4, BA.5, and BA.2.75 when compared to relatively older BA.1 and BA.2 lineages (**Fig. 1c–e, Extended Data Fig. 1a**). For instance, the R346X and K444X substitutions in BA.4 and BA.5 and the R346X and F486X substitutions in BA.2.75 showed substantially higher substitution frequencies compared to those in the other lineages (**Fig. 1c**). The substitution frequencies at R346 and K444 in BA.5 were approximately 10.4- and 9.4-times higher than those in BA.2, respectively (**Fig. 1e and Supplementary Table 1**).

**Fig. 1.**
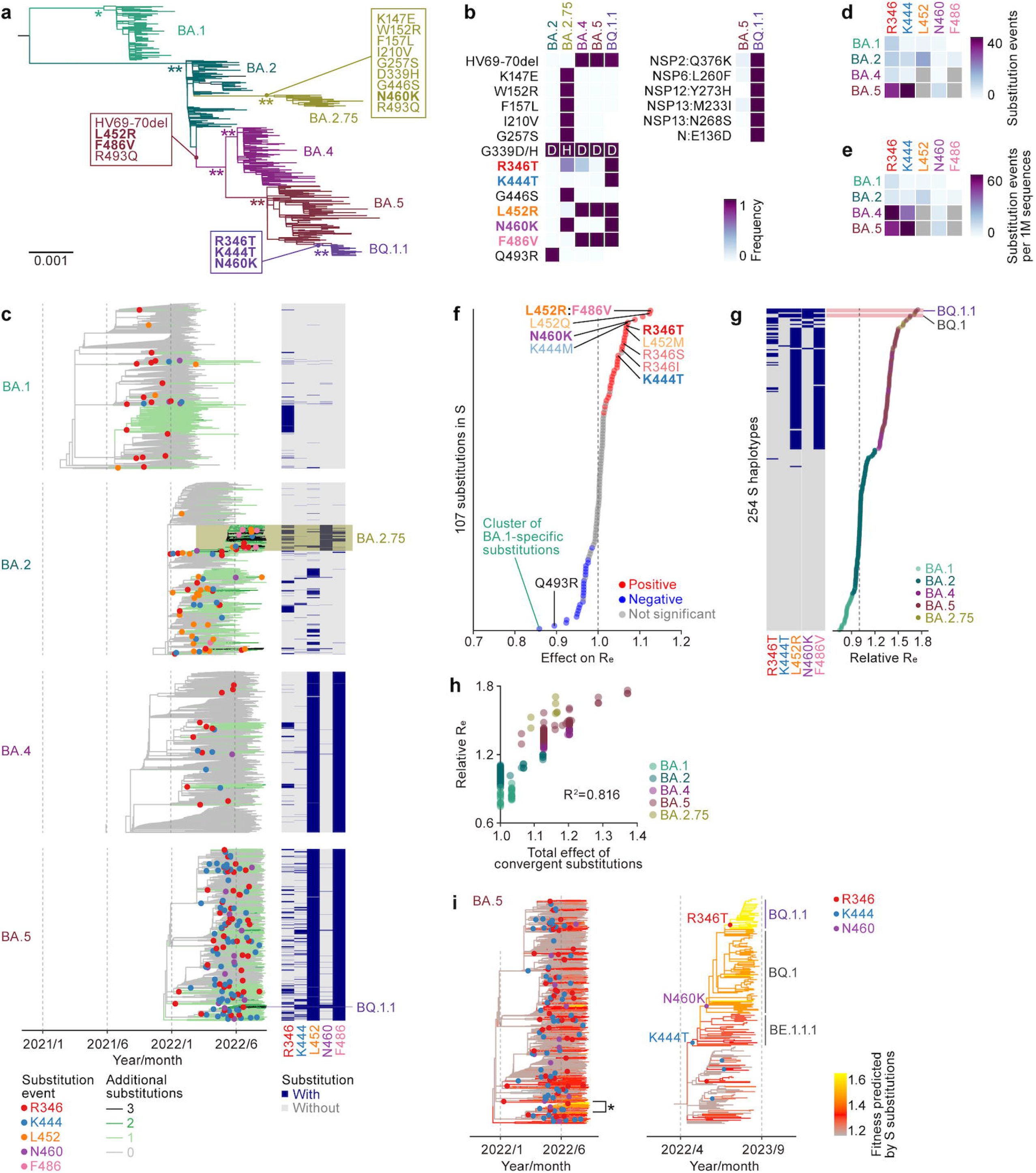
Convergent evolution of Omicron lineages. **a,** A maximum likelihood (ML) tree of the Omicron lineages. The tree was rooted using an outgroup sequence (B.1.1). The S substitutions acquired by BA.4/BA.5, BA.2.75, and BQ.1.1 are indicated in the panel, and the five convergent substitutions are indicated in bold. Note that R493Q is a reversion. Bootstrap values, *, ≥ 0.85; **, ≥ 0.9. **b,** Left, amino acid differences in the S proteins of Omicron lineages. The five convergent substitutions are indicated in bold. Right, amino acid differences in the non-S proteins between BA.5 and BQ.1.1. **c,** Left, time-calibrated ML trees for BA.1, BA.2, BA.4, and BA.5. The trees for BA.2 and BA.5 include BA.2.75 and BQ.1.1 lineages, respectively. Dots indicate estimated substitution events at the convergent sites. Branch color indicates the estimated number of additional substitutions at the convergent sites compared to the most recent common ancestor of each lineage. Right, the substitution profile at the convergent sites. **d and e,** The number of substitution events at the convergent sites detected. Raw counts (**d**) and counts per 1 million (1M) analyzed sequences (**d**) are shown. Note that L452 and F486 in BA.4/5 are indicated in gray because BA.4/5 harbors the L452R and F486V substitutions. **f,** Effect size of each S substitution on R_e_ estimated by a hierarchal Bayesian model. Posterior mean value is shown. A group of highly co-occurred substitutions (e.g., L452R and F486V) was treated as substitution clusters. The red and blue dots indicate the substitutions with significant positive and negative effects, respectively. Representative substitutions are annotated. **g,** Relative R_e_ value for a viral group represented by each S haplotype, assuming a fixed generation time of 2.1-day. Posterior mean value is shown. The R_e_ of the major S haplotype in BA.2 is set at 1. The substitution profile at the five convergent sites is shown in the left. **h,** Comparison between relative R_e_ and the total effect of substitutions at the convergent sites on R_e_. Dot indicates a viral group represented by a S haplotype. Dots are colored according to the major classification of PANGO lineage. **i,** Change in viral fitness during BA.5 diversification. The lineages indicated with an asterisk, which includes BQ.1.1, are zoomed in the right panel.

### Amino acid substitutions at the convergent sites increase viral fitness

We next hypothesized that the substitutions at these five convergent sites conferred certain selective advantages during the evolution of Omicron. To test this hypothesis, we modeled the relationship between viral epidemic dynamics and the substitutions in S protein and estimated the effects of all S substitutions on viral fitness [i.e., relative effective reproduction number (R_e_)]. We classified the viral sequences of Omicron according to the combination of amino acid substitutions (referred to as “S haplotype”). Subsequently, we established a Bayesian hierarchal model, which represents the epidemic dynamics of the S haplotype-based viral groups according to relative R_e_ represented by a linear combination of the effect of S substitutions (see **Methods**). This model can simultaneously estimate i) the effect of each S substitution on R_e_ and ii) the relative R_e_ of a viral group represented by each S haplotype. We analyzed the dataset for 375,121 Omicron sequences collected in the United Kingdom (UK) from March 1^st^, 2022, to October 15^th^, 2022, which includes 254 S haplotypes according to the pattern of 107 substitutions in S protein [or substitution clusters (**Extended Data Fig. 1b**)]. We first investigated the effects of S substitution on R_e_ (**Fig. 1f, Extended Data Fig. 1c and Supplementary Table 2**). As substitutions with negative effects on Re, the cluster of substitutions specific to BA.1 – the earliest Omicron lineage with the lowest R_e_ overall – and Q493R – acquired once in the common ancestor of Omicron but subsequently lost in BA.5 and BA.2.75 – were identified (**Fig. 1a,f, Extended Data Fig. 1c and Supplementary Table 2**). On the other hand, as substitutions with positive effects on Re, substitutions at the five convergent sites were identified (**Fig. 1f, Extended Data Fig. 1c and Supplementary Table 2**). Particularly, i) the L452R and F486V substitutions, acquired by the common ancestor of BA.4 and BA.5, ii) L452Q, acquired by BA.2.12.1^15^, and iii) N460K, acquired by BA.2.75 and BQ.1.1 independently, showed higher positive effects (**Fig. 1f, Extended Data Fig. 1c and Supplementary Table 2**). Next, we investigated relative R_e_ values for viral groups represented by respective S haplotypes (**Fig. 1g, Extended Data Fig. 1d and Supplementary Table 3**). We found that S haplotypes with substitutions at the convergent sites, particularly with R346T, K444T, L452R, N460K, and F486V, tended to show higher R_e_ values (**Fig. 1g**). Notably, the S haplotype corresponding to BQ.1.1, harboring all five aforementioned convergent substitutions, showed the highest R_e_ value, followed by S haplotypes corresponding to BQ.1, harboring all substitutions apart from R346T (**Fig. 1g and Supplementary Table 3**).

To quantify the impact of substitutions at the convergent sites for viral fitness, we inferred the proportion of the variation of R_e_ that can be explained by these substitutions in the Omicron lineages. We first calculated the total effect of substitutions at the convergent sites for each S haplotype. Subsequently, we compared this quantity with the relative R_e_ value for each S haplotype (**Fig. 1h**). These two quantities were strongly correlated (R^2^=0.816), suggesting that a larger proportion (81.6%) of the variation of R_e_ in the Omicron lineages can be explained by substitutions at the convergent sites under our model.

Using the Bayesian hierarchal model above, we can predict relative viral fitness for arbitrary S sequences only based on the profile of S substitutions (see **Methods**). Utilizing this property of this model, we next inferred the evolutionary change in viral fitness during BA.5 diversification. We reconstructed the ancestral profile of S substitutions for each node in the BA.5 tree. Subsequently, we predicted relative viral fitness for each node according to the reconstructed S substitution profile using the model above (**Fig. 1i**). This analysis suggested that the viral fitness was independently elevated in multiple lineages during BA.5 diversification, coupled with the substitution events at the convergent sites (**Fig. 1i, left**). Finally, we inferred the evolutionary changes in viral fitness specific to the emergence of BQ.1.1 and revealed that the ancestral lineage of BQ.1.1 has acquired the K444T, N460K, and R346T substitutions in this order (**Fig. 1i, right**)^20^. Importantly, our analysis predicted that the ancestral lineage of BQ.1.1 has increased its viral fitness step-by-step consistently with the acquisitions of these substitutions (**Fig. 1i, right**). Taken together, our results suggest that the sublineages descending from BA.5, including BQ.1.1, convergently increased viral fitness by consecutively acquiring substitutions at the R346, N460, and K444 residues.

### Immune resistance of BQ.1.1

It has been recently reported that BQ.1.1 exhibits profound resistance to all therapeutic monoclonal antibodies currently approved by the Food and Drug Administration (FDA) in the United States^6,21^. Also, some substitutions detected in BQ.1.1, such as R346T and K444T, contribute to the resistance to 3-dose treatment of an inactivated vaccine (CoronaVac) and breakthrough infections or prior Omicron subvariants (including BA.1, BA.2 and BA.5) after CoronaVac vaccination^6^. However, the immune resistance of BQ.1.1 to breakthrough infections or prior Omicron subvariants after mRNA vaccine treatment remains unaddressed. To experimentally investigate the virological features of BQ.1.1, we first evaluated the immune resistance of BQ.1.1 using HIV-1-based pseudoviruses. Consistent with our recent study^15^, BA.5 (2.5-fold) and BQ.1.1 (6.9-fold) were significantly more resistant to breakthrough BA.2 infection sera than BA.2 (**Fig. 2a**). Additionally, BQ.1.1 exhibited more profound resistance to breakthrough BA.2 infection sera than BA.5 (2.7-fold, p=0.0076) (**Fig. 2a**). As shown in **Fig. 1a**, the BQ.1.1 S harbors three additional substitutions in the BA.5 S: R346T, K444T and N460K. To determine the responsible substitution(s) for immune resistance of BQ.1.1 to breakthrough BA.2 infection sera, we prepared the BA.5 derivatives bearing either of these three substitutions. However, any BA.5-based derivatives prepared did not exhibit resistance to breakthrough BA.2 infection sera compared to BA.5 (**Fig. 2a**), suggesting that multiple substitutions cooperatively contribute to the immune resistance of BQ.1.1 to breakthrough BA.2 infection sera. In the case of breakthrough BA.5 infection sera, BQ.1.1 was significantly (5.6-fold) more resistant to breakthrough BA.5 infection sera than BA.5 (p<0.0001) (**Fig. 2b**). Importantly, the breakthrough BA.5 infection sera obtained from six individuals (five breakthrough infection cases after 3-dose vaccination and a breakthrough infection case after 2-dose vaccination) did not exhibit antiviral effect against BQ.1.1 in this experimental setup. We then assessed the determinant substitutions to be resistant to breakthrough BA.5 infection sera. The N460K substitution conferred significant resistance to breakthrough BA.5 infection sera (1.6-fold, p=0.016), while the other two substitutions did not affect the immune resistance to breakthrough BA.5 infection sera (**Fig. 2b**). When compared to the immune resistance of BQ.1.1 to breakthrough BA.5 infection sera (5.6-fold), the resistance acquired by the N460K substitution (1.6-fold) is relatively less robust (**Fig. 2b**). Therefore, similar to breakthrough BA.2 infection sera, our results suggest that the immune resistance of BQ.1.1 to breakthrough BA.2 infection sera is attributed by multiple substitutions in the RBD of BQ.1.1 S.

**Fig. 2.**
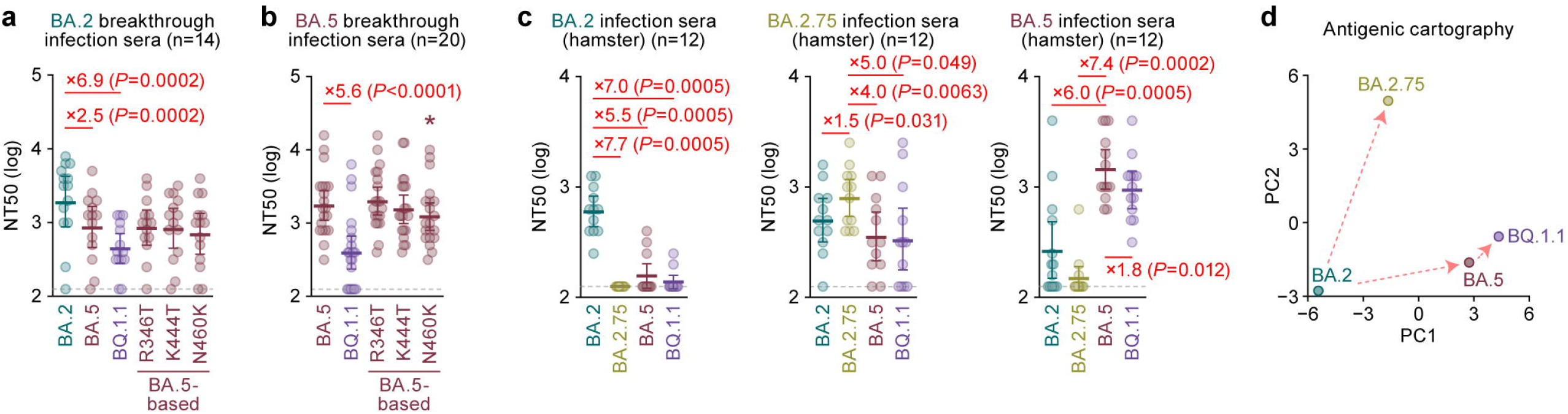
Immune resistance of BQ.1.1. Neutralization assays were performed with pseudoviruses harboring the S proteins of BA.2, BA.2.75, BA.5 and BQ.1.1. The BA.5 S-based derivatives are included in **a and b**. The following sera were used. **a,b,**Convalescent sera from fully vaccinated individuals who had been infected with BA.2 after full vaccination (9 2-dose vaccinated and 5 3-dose vaccinated. 14 donors in total) (**a**), and BA.5 after full vaccination (2 2-dose vaccinated donors, 17 3-dose vaccinated donors and 1 4-dose vaccinated donors. 20 donors in total) (**b**). **c,** Sera from hamsters infected with BA.2 (12 hamsters; left), BA.2.75 (12 hamsters; middle), and BA.5 (12 hamsters; right). **d,** Principal component analysis to representing the antigenicity of the S proteins. The analysis is based on the results of neutralization assays using hamster sera (**Fig. 3c**). Assays for each serum sample were performed in triplicate to determine the 50% neutralization titer (NT_50_). Each dot represents one NT_50_ value, and the geometric mean and 95% CI are shown. Statistically significant differences were determined by two-sided Wilcoxon signed-rank tests. The *P* values versus BA.2 (**a and c, left**) BA.2.75 (**c, middle**) or BA.5 (**b and c, right**) are indicated in the panels. For the BA.5 derivatives (**a and b**), statistically significant differences (*P* < 0.05) versus BA.5 are indicated with asterisks. The horizontal dashed line indicates the detection limit (120-fold). Information on the convalescent donors is summarized in **Supplementary Table 4**.

To further address the difference in antigenicity among Omicron subvariants including BQ.1.1, we used the sera obtained from infected hamsters at 16 days postinfection (d.p.i.). Consistent with our previous studies^2,15^, the hamster sera infected with BA.2, BA.5 or BA.2.75 were most efficiently antiviral against the variant of virus infected, while these antisera were less or no cross-reactive against the other variants (**Fig. 2c**). In the case of BA.5 infection sera, BQ.1.1 was 1.8-fold more resistant to than BA.5 (**Fig. 2c**). To depict the difference of antigenicity among BA.2, BA.5, BA.2.75 and BQ.1.1, we analyzed the neutralization dataset using hamster sera (**Fig. 2c**). As shown in **Fig. 2d**, the cross-reactivity of each Omicron subvariant was well correlated to their phylogenetic relationship (**Fig. 1a**), and the antigenicity of BQ.1.1 is relatively more similar to BA.5 than BA.2 and BA.2.75. Nevertheless, our data show that BQ.1.1 exhibits a profound resistance to the humoral immunity induced by BA.5 breakthrough infection (**Fig. 2b**). These observations suggest that the three substitutions in BQ.1.1 S are critical and specific to evade the BA.5 infection-induced herd immunity in the human population.

### ACE2 binding affinity of BQ.1.1 S

We then evaluated the features of BQ.1.1 S that potentially affect viral infection and replication. Yeast surface display assay^2,15,18,19,22–24^ showed that the K_D_ value of BQ.1.1 S RBD (0.66±0.11) to human ACE2 molecule is significantly lower than that of parental BA.5 S RBD (1.08±0.16) (**Fig. 3a**), suggesting that BQ.1.1 increased the binding affinity to human ACE2 during evolution from BA.5. To determine the responsible substitutions in the BQ.1.1 S that enhance ACE2 binding affinity, we prepared the RBDs of BA.2 and BA.5 S proteins that possess a BQ. 1.1-specific substitution compared to parental BA.5 (i.e., R346T, K444T and N460K). Consistent with our recent study^2^, the N460K substitution significantly increased the binding affinity of the S proteins of BA.2 and BA.5 to human ACE2 (**Fig. 3a**). On the other hand, the K444 substitution significantly decreased ACE2 binding affinity regardless of the S backbone (**Fig. 3a**). The R346T substitution increased the ACE2 binding affinity of BA.2 S RBD but not that of BA.5 S RBD (**Fig. 3a**). The *in vitro* observations using yeasts (**Fig. 3a**) were then verified by using an HIV-1-based pseudovirus system. As shown in **Fig. 3b**, the infectivity of BQ.1.1 pseudovirus was significantly higher than those of BA.2 (17-fold) and BA.5 (3.2-fold) pseudoviruses. In our recent study^15^, at least three mutations detected in the BA.5 S (compared to the BA.2 S), HV69-70del, L452R and F486V contribute to the increase of pseudovirus infectivity. When we particularly focus on the three additional mutations detected in the BQ.1.1 S compared to the BA.5 S, R346T, K444T and N460K, we found that R346T and N460K but not K444T significantly increase pseudovirus infectivity, and it is independent of the S backbone (**Fig. 3b**). To assess the association of TMPRSS2 usage with the increased pseudovirus infectivity of BQ.1.1, we used HEK293-ACE2/TMPRSS2 cells and HEK293-ACE2 cells, on which endogenous surface TMPRSS2 is undetectable^18^ as target cells. As shown in **Fig. 3c**, the infectivity of BQ.1.1 pseudovirus was not increased by TMPRSS2 expression, suggesting that TMPRSS2 is not associated with an increase in the infectivity of BQ.1.1 pseudovirus.

**Fig. 3.**
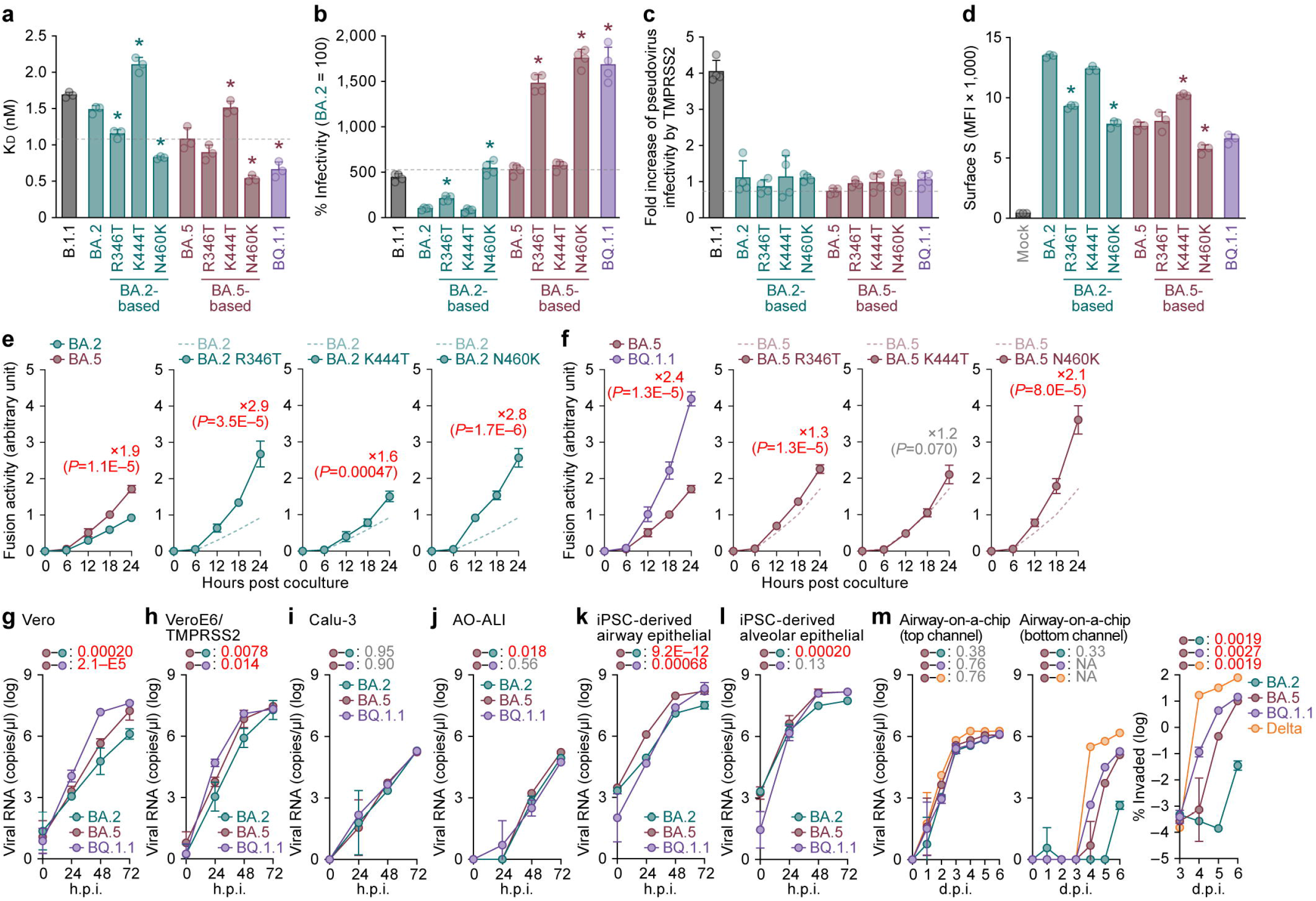
Virological characteristics of BQ.1.1 *in vitro*. **a,** Binding affinity of the RBD of SARS-CoV-2 S protein to ACE2 by yeast surface display. The K_D_ value indicating the binding affinity of the RBD of the SARS-CoV-2 S protein to soluble ACE2 when expressed on yeast is shown. **b,** Pseudovirus assay. HOS-ACE2-TMPRSS2 cells were infected with pseudoviruses bearing each S protein. The amount of input virus was normalized based on the amount of HIV-1 p24 capsid protein. The percent infectivity compared to that of the virus pseudotyped with the BA.2 S protein are shown. **c,** Fold increase in pseudovirus infectivity based on TMPRSS2 expression. **d–f,** S-based fusion assay. **d,** S protein expression on the cell surface. The summarized data are shown. **e,f,** S-based fusion assay in Calu-3 cells. The recorded fusion activity (arbitrary units) is shown. The dashed green line (**e**) and the dashed brown line (**f**) indicate the results of BA.2 and BA.5, respectively. The red number in each panel indicates the fold difference between BA.2 (**e**) or BA.5 (**f**) and the derivative tested at 24 h post coculture. **g–l,** Growth kinetics of BQ.1.1. Clinical isolates of BA.2, BA.5, BQ.1.1 and Delta (only in **m**) were inoculated into Vero cells (**g**), VeroE6/TMPRSS2 cells (**h**), Calu-3 cells (**i**), AO-ALI (**j**), iPSC-derived airway epithelial cells (**k**), iPSC-derived alveolar epithelial cells (**l**) and an airway-on-a-chip system (**m**). The copy numbers of viral RNA in the culture supernatant (**g–i**), the apical sides of cultures (**j–l**), and the top (**m, left**) and bottom (**m, middle**) channels of an airway-on-a-chip were routinely quantified by RT–qPCR. In **m, right**, the percentage of viral RNA load in the bottom channel per top channel during 3–6 d.p.i. (i.e., % invaded virus from the top channel to the bottom channel) is shown. Assays were performed in triplicate (**a,d,m**) or quadruplicate (**b,c,e–l**). The presented data are expressed as the average ± SD (**a–f**) or SEM (**g–m**). In **a–d**, each dot indicates the result of an individual replicate. In **a–c**, the dashed horizontal lines indicate the value of BA.5. In **a–d**, statistically significant differences (*, *P* < 0.05) versus each parental S and those between BA.5 and BQ.1.1 were determined by two-sided Student’s *t* tests. In **e–m**, statistically significant differences versus BA.2 (**e**) and BA.5 (**f–m**) across timepoints were determined by multiple regression. The FWERs calculated using the Holm method are indicated in the figures. NA, not applicable.

### Fusogenicity of BQ.1.1 S

The fusogenicity of BQ.1.1 S was measured by the SARS-CoV-2 S-based fusion assay^2,15–19,25^ using Calu-3 cells. Surface expression level of the BQ.1.1 S was significantly lower than that of BA.2, but the BQ.1.1 and BA.5 expression level were comparable (**Fig. 3d**). In the BA.2 S derivatives, R346T and N460K significantly decreased surface expression (**Fig. 3d**). In the BA.5 S derivatives, N460K significantly decreased surface expression, while K444T increased it (**Fig. 3d**). The fusogenicity of BA.5 S is greater than that of BA.2 S (**Fig. 3e**), which is consistent with our recent studies^2,15^. More importantly, the BQ.1.1 S was significantly more fusogenic than the BA.5 S (**Fig. 3f**). Additional experiments using the S derivatives based on BA.2 and BA.5 showed that the R346T and N460K substitutions significantly increased the S-mediated fusogenicity independently of the S backbone (**Fig. 3e,f**). Together with our recent studies^2^, the N460K substitution, which is detected in BA.2.75, increased ACE2 binding affinity (i.e., decrease of the K_D_ value) (**Fig. 3a**), increased pseudovirus infectivity (**Fig. 3b**) and the S-mediated fusogenicity (**Fig. 3e,f**). Interestingly, the R346T substitution also significantly increased ACE2 binding affinity and the S-based fusogenicity, while the K444T substitution negatively affected these experimental parameters (**Fig. 3b–f**). These results suggest that, compared to BA.5, the virological features of BQ.1.1 S, including increased ACE2 binding affinity, pseudovirus infectivity and fusogenicity, are attributed to the R346T and N460K substitutions.

### Growth kinetics of BQ.1.1 *in vitro*

To investigate the growth kinetics of BQ.1.1 in *in vitro* cell culture systems, we inoculated clinical isolates of BA.2^18^, BA.5^2^ and BQ.1.1 into multiple cell cultures. The growth of BQ.1.1 in Vero cells (**Fig. 3g**) and VeroE6/TMPRSS2 cells (**Fig. 3h**) was significantly greater than that of BA.5, and the growth of BQ.1.1 and BA.5 was comparable in Calu-3 cells (**Fig. 3i**), human airway organoid-derived air-liquid interface (AO-ALI) system (**Fig. 3j**), and human induced pluripotent stem cell (iPSC)-derived alveolar epithelial cells (**Fig. 3l**). However, the growth of BQ.1.1 in iPSC-derived airway epithelial cells was significantly lower than that of BA.5 (**Fig. 3k**).

To evaluate the impact of BQ.1.1 infection on the airway epithelial and endothelial barriers, we used an airway-on-a-chip system^2,26^. By measuring the amount of virus that invades from the top channel (**Fig. 3m, left**) to the bottom channel (**Fig. 3m, middle**), we are able to evaluate the ability of viruses to disrupt the airway epithelial and endothelial barriers. Notably, the percentage of virus that invaded the bottom channel of BQ.1.1-infected airway-on-chip was significantly higher than that of BA.5-infected airway-on-chip (**Fig. 3m, right**). Together with the findings of S-based fusion assay (**Fig. 3f**), these results suggest that BQ.1.1 has higher fusogenic than BA.5.

### Virological characteristics of BQ.1.1 *in vivo*

To investigate the virological features of BQ.1.1 *in vivo,* we inoculated clinical isolates of Delta^16^, BA.5^2^, and BQ.1.1. Consistent with our previous studies^2,16^, Delta infection resulted in weight loss (**Fig. 4a, left**). On the other hand, the body weights of BA.5- and BQ.1.1-infected hamsters did not increase compared with the negative control and relatively similar (**Fig. 4a, left**). We then analyzed the pulmonary function of infected hamsters as reflected by two parameters, enhanced pause (Penh) and the ratio of time to peak expiratory flow relative to the total expiratory time (Rpef). Among the four groups, Delta infection resulted in significant differences in these two respiratory parameters compared to BA.5 (**Fig. 4a, middle and right**), suggesting that Delta is more pathogenic than BA.5. In contrast, the Penh value of BQ.1.1-infected hamsters was significantly lower than that of BA.5-infected hamsters, and the Rpef value of BQ.1.1-infected hamsters was significantly higher than that of BA.5-infected hamsters (**Fig. 4a, middle and right**). These observations suggest that the pathogenicity of BQ. 1. 1 is similar to or even less than that of BA.5.

**Fig. 4.**
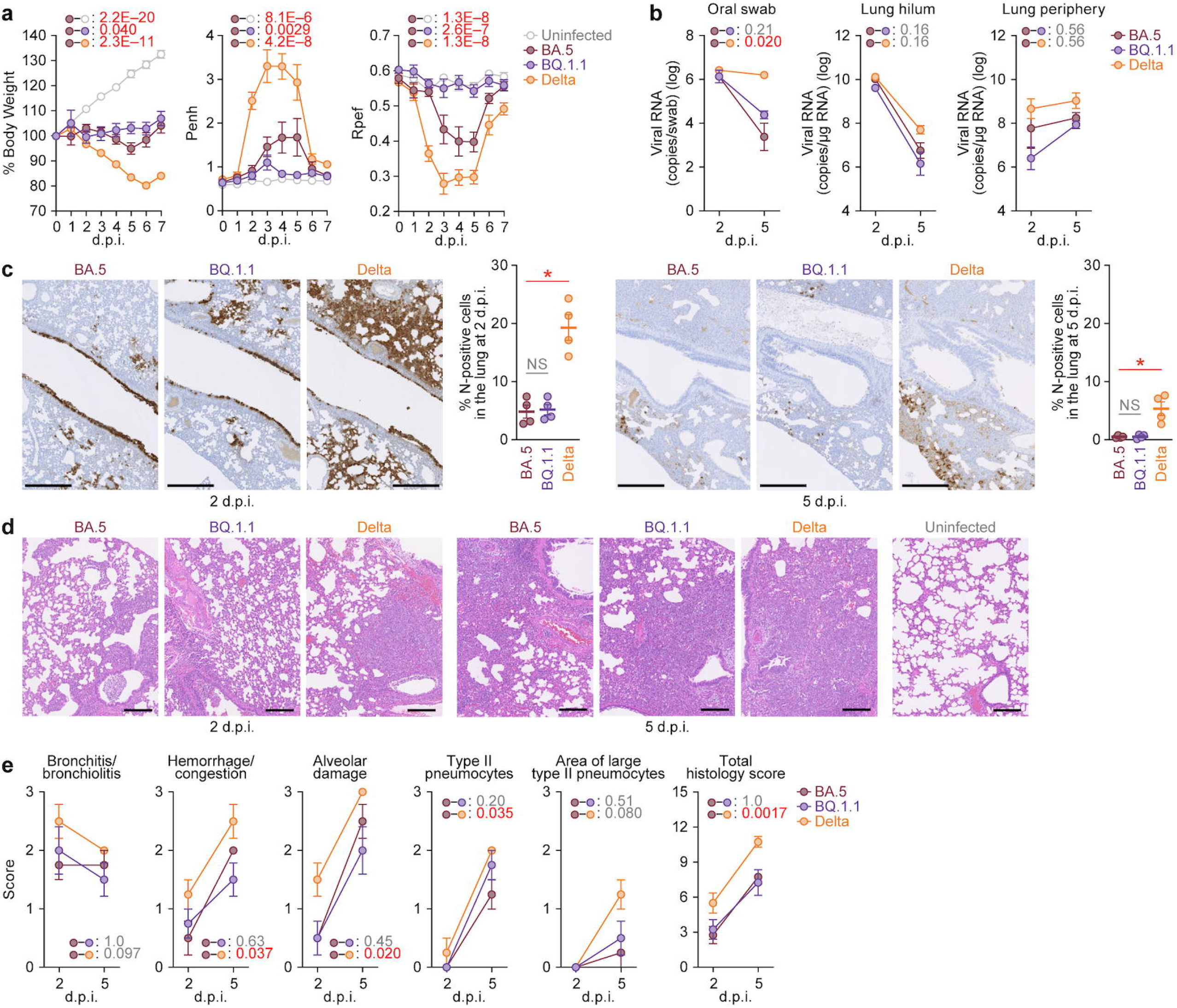
Virological characteristics of BQ.1.1 *in vivo*. Syrian hamsters were intranasally inoculated with BA.5, BQ.1.1 and Delta. Six hamsters of the same age were intranasally inoculated with saline (uninfected). Six hamsters per group were used to routinely measure the respective parameters (**a**). Four hamsters per group were euthanized at 2 and 5 d.p.i. and used for virological and pathological analysis (**b–e**). **a,** Body weight, Penh, and Rpef values of infected hamsters (n = 6 per infection group). **b,** (Left) Viral RNA loads in the oral swab (n=6 per infection group). (Middle and right) Viral RNA loads in the lung hilum (middle) and lung periphery (right) of infected hamsters (n=4 per infection group). **c,** IHC of the viral N protein in the lungs at 2 d.p.i. (left) and 5 d.p.i. (right) of infected hamsters. Representative figures (N-positive cells are shown in brown) and the percentage of N-positive cells in whole lung lobes (n=4 per infection group) are respectively shown. The raw data are shown in **Extended Data Fig. 2b**. **d, e,** H&E staining of the lungs of infected hamsters. Representative figures are shown in **d**. Uninfected lung alveolar space and bronchioles are also shown. **e,**Histopathological scoring of lung lesions (n=4 per infection group). Representative pathological features are reported in our previous studies^27^. In **a–c,e**, data are presented as the average ± SEM. In **c**, each dot indicates the result of an individual hamster. In **a,b,e**, statistically significant differences between BA.5 and other variants across timepoints were determined by multiple regression. In **a**, the 0 d.p.i. data were excluded from the analyses. The FWERs calculated using the Holm method are indicated in the figures. In **c**, the statistically significant differences between BA.5 and other variants were determined by a two-sided Mann–Whitney *U* test. In **c and d**, each panel shows a representative result from an individual infected hamster. Scale bars, 500 μm (**c**); 200 μm (**d**).

To evaluate the viral spread in infected hamsters, we routinely measured the viral RNA load in the oral swab. Although the viral RNA loads of the hamsters infected with Delta were significantly higher than those infected with BA.5, there was no statistical difference between BQ.1.1 and BA.5 (**Fig. 4b, left**). To address the possibility that BQ.1.1 more efficiently spreads in the respiratory tissues, we collected the lungs of infected hamsters at 2 and 5 d.p.i., and the collected tissues were separated into the hilum and periphery regions. However, the viral RNA loads in both lung hilum and periphery of BA.5-infected hamsters were comparable to those of Delta- and BQ.1.1-infected hamsters (**Fig. 4b, middle and right**), suggesting that the spreading efficacy of BQ.1.1 in the lungs is comparable to BA.5. To further investigate the viral spread in the respiratory tissues of infected hamsters, we performed immunohistochemical (IHC) analysis targeting viral nucleocapsid (N) protein. Similar to our previous studies^15,17,18^, epithelial cells in the upper tracheae of infected hamsters were sporadically positive for viral N protein at 2 d.p.i., but there were no significant differences among the three viruses, including BQ.1.1 (**Extended Data Fig. 2a**), although tracheal inflammation tended to remain in BQ.1.1-infected hamsters. In the alveolar space around the bronchi/bronchioles at 2 d.p.i., N-positive cells were detected in Delta-infected hamsters (**Fig. 4c, left and Extended Data Fig. 2b**). In contrast, the percentage of N-positive cells in the lungs of BQ.1.1- and BA.5-infected hamsters were relatively low and comparable (**Fig. 4c, left and Extended Data Fig. 2b**). At 5 d.p.i., N-positive cells were detected in the peripheral alveolar space in Delta-infected hamsters, while the N-positive areas of BQ.1.1- and BA.5-infected hamsters were sporadic and faintly detectable (**Fig. 4c, right and Extended Data Fig. 2b**). These data suggest that the spreading efficiency of BQ.1.1 in the lungs of infected hamsters is comparable to that of BA.5.

### Intrinsic pathogenicity of BQ.1.1

To investigate the intrinsic pathogenicity of BQ.1.1, we analyzed the formalin-fixed right lungs of infected hamsters at 2 and 5 d.p.i. by carefully identifying the four lobules and main bronchus and lobar bronchi sectioning each lobe along with the bronchial branches (**Fig. 4d**). Histopathological scoring was performed according to the criteria described in our previous studies^27^. Consistent with our previous studies^2,16,17^, all five histological parameters as well as the total score of the Delta-infected hamsters were significantly greater than those of the BA.5-infected hamsters (**Fig. 4e**). When we compared the histopathological scores of Omicron subvariants, total histopathological scores were comparable between BQ.1.1-infected hamsters and BA.5-infected hamsters with some enhancement of bronchitis/bronchiolitis at 2 d.p.i. and presence of type II pneumocytes at 5 d.p.i. of BQ.1.1 (**Fig. 4e**). Altogether, these histopathological analyses suggest that the intrinsic pathogenicity of BQ.1.1 is lower than that of Delta and comparable to that of BA.5.

## Discussion

In the present study, we highlighted the amino acid substitutions that convergently and frequently occurred in the residues R346, K444, L452, N460, and F486 of the S proteins of relatively younger Omicron lineages, such as BA.4, BA.5, and BA.2.75 (**Fig. 1a,c**). Our modeling analysis suggested that these five frequent substitutions at the convergent sites increase viral fitness, R_e_ (**Fig. 1f**), and a larger proportion (81.6%) of the R_e_ variation within Omicron can be explained by substitutions in the convergent sites (**Fig. 1h**). Intriguingly, the viral groups harboring the substitutions in convergent sites showed higher Re, and BQ.1.1, which harbors all convergent substitutions, showed the highest R_e_ among viral groups investigated (**Fig. 1g**). Moreover, the reconstruction of ancestral viral fitness suggests that the ancestral lineage of BQ.1.1 has increased viral fitness by acquiring substitutions at the convergent sites in a stepwise manner (**Fig. 1i**). Altogether, our integrated approach of phylogenetic and epidemic dynamics modeling analyses provides insights into the evolutionary rules underlying the outstanding convergent evolution observed in Omicron subvariants.

Our data provide two possibilities that explain the accumulated amino acid substitutions at the convergent sites in relatively younger Omicron lineages, such as BA.4, BA.5 and BA.2.75. One possibility is the epistasis among amino acid substitutions: the fitness of a substitution differs dependent on the presence of the other substitutions and/or the backbone sequence (similar to how the original Omicron genotype likely emerged^28^). In our previous studies, the L452R substitution in the BA.4/5 S^15^ and the N460K substitution in the BA.2.75 S^2^ increase their binding ability to human ACE2. More importantly, we showed that these substitutions can compensate for the negative effects of the other substitutions that contribute to evasion from antiviral humoral immunity but decrease ACE2 binding ability: for BA.4/5, L452R compensates for the attenuated ACE2 binding affinity by the F486V substitution^4,7–9^, while for BA.2.75, N460K compensates for the attenuated ACE2 binding affinity by the G446S substitution^2,7,11^. Acquiring substitutions that potentially increase ACE2 binding ability, such as L452R and N460K, can be one of the factors that increases substitution frequency at the convergent sites in BA.4, BA.5, and BA.2.75. The other possibility is that the effect of substitutions on viral fitness can be changed over time due to changes in immune selective pressures in the human population by vaccinations and/or natural infections with a variety of SARS-CoV-2 variants. The substitutions at R346, K444, and F486 closely associate with the escape from antiviral humoral immunity and monoclonal antibodies^5,6,15^. In fact, here we demonstrated that the antigenicity of BQ.1.1 is different from that of parental BA.5 and BQ.1.1 is more robustly resistant to the antiviral humoral immunity induced by BA.2 and BA.5 breakthrough infections than BA.5. Therefore, it is reasonable to assume that the effect of substitutions at these sites on viral fitness likely differs depending on the immune status in the human population. These two possibilities are not mutually exclusive, and these two factors could contribute to the accelerated substitutions at the convergent sites.

Based on our experimental results as well as recent reports, it is convincing that BQ.1.1 is one of the variants exhibiting profound resistance to the antiviral humoral immunity induced by breakthrough infections of BA.2 and BA.5 as well as therapeutic monoclonal antibodies^6,21,29^. Additionally, our experiments *in vitro* showed that the BQ.1.1 S exhibits higher affinity to human ACE2 and greater fusogenicity than BA.5, and these abilities are conferred by two substitutions, R346T and N460K. As expected, because BQ.1.1 bears two substitutions, L452R^15^ and N460K^2^, both of which augment ACE2 binding affinity, BQ.1.1 has evolved to augment its fusogenicity. Moreover, the R346T substitution, while contributing to enhanced ACE2 affinity and augmented fusogenicity, also concomitantly contributes, perhaps unexpectedly, to immune evasion^6,21^.

Our previous studies focusing on Delta^16^, Omicron BA.1^17^, BA.5^15^, and BA.2.75^2^, showed that the intrinsic pathogenicity of SARS-CoV-2 variants is closely related to the fusogenicity of viral S proteins. Therefore, the observations showing the higher fusogenicity of BQ.1.1 S than the BA.5 S based on the S-based fusion assay (**Fig. 3f**) and the experiments using airway-on-a-chip (**Fig. 3m**) suggested the increased intrinsic pathogenicity of BQ.1.1 when compared to BA.5. However, it was unexpected that the intrinsic pathogenicity of BQ.1.1 in a hamster model is comparable or even lower than that of BA.5 (**Fig. 4**). This discrepancy between viral fusogenicity and viral intrinsic pathogenicity is reminiscent of the previous two studies on Omicron BA.2 variant. We first showed that the BA.2 S is more fusogenic than the BA.1 S^18^. We then artificially generated a BA.2 S-bearing recombinant SARS-CoV-2, in which the non-S region of viral genome is derived from ancestral SARS-CoV-2 and demonstrated that the BA.2 S-bearing virus is more pathogenic than the BA.1 S-bearing virus in hamsters^18^. On the other hand, Uraki et al. showed that the intrinsic pathogenicity of clinical BA.2 isolates is comparable to clinical BA.1 isolates^30^. Because the difference between our study^18^ and others^30^ is the viral genome sequence in the non-S region, it is suggested that the BA.2 S bears the potential to exhibit augmented pathogenicity when compared to the BA.1 S, whereas the mutations in non-S region of BA.2 genome potentially attenuate viral pathogenicity. In fact, we found there are at least six substitutions in the non-S region of BQ.1.1 when compared to that of BA.5. Therefore, it would be conceivable to assume that there are factors potentially modulate viral intrinsic pathogenicity other than S protein.

There are two evolutionary scenarios that possibly explain the discrepancy between viral fusogenicity and intrinsic pathogenicity observed in BQ.1.1. One scenario is that BQ.1.1 acquired mutation(s) in the non-S region of viral genome that can attenuate viral pathogenicity and cancel the pathogenicity elevated by the higher fusogenicity compared with the parental BA.5. This is reminiscent to the observations in the two BA.2 studies as mentioned above^18,30^. This scenario also provides a possibility that the increased fusogenicity of viral S protein can reduce viral fitness in the human population because greater fusogenicity can result in elevating pathogenicity. Another scenario is brought from a theoretical study by Sasaki, Lion, and Boots^31^. This study provides a possibility that antigenic escape can augment viral pathogenicity^31^. Since we demonstrated that at least two descendants of BA.2, BA.5^15^, and BA.2.75^2^, increased their intrinsic pathogenicity, this theory may fit the evolution of Omicron. More importantly, this theory also predicts that there is a limitation to increase viral pathogenicity^31^. Together with our observations, it might be possible to assume that the pathogenicity of Omicron lineage already reaches a plateau.

The emergence of SARS-CoV-2 variants with increased intrinsic pathogenicity may not be so critical for the immunized segment of the population. However, the variants bearing greater intrinsic pathogenicity can be a meaningful risk for people who do not have anti-SARS-CoV-2 immunity, most conspicuously the unvaccinated population, including children. Therefore, continued in-depth viral genomic surveillance and real-time evaluation of the risk of newly emerging SARS-CoV-2 variants should be crucial.

## Author Contributions

Keiya Uriu, Sayaka Deguchi, Hesham Nasser, Maya Shofa, MST Monira Begum, Shigeru Fujita, Akatsuki Saito, Kazuo Takayama and Terumasa Ikeda performed cell culture experiments.

Rigel Suzuki, Yukari Itakura, Tomokazu Tamura, Izumi Kida, Kumiko Yoshimatsu, Hayato Ito, Naganori Nao, Keita Matsuno and Takasuke Fukuhara performed animal experiments.

Lei Wang, Masumi Tsuda, Yoshitaka Oda and Shinya Tanaka performed histopathological analysis.

Jiri Zahradnik and Gideon Schreiber performed yeast surface display assay. Sayaka Deguchi and Kazuo Takayama prepared AO-ALI and airway-on-a-chip systems.

Yuki Yamamoto and Tetsuharu Nagamoto performed generation and provision of human iPSC-derived airway and alveolar epithelial cells.

Hiroyuki Asakura, Mami Nagashima, Kenji Sadamasu and Kazuhisa Yoshimura performed viral genome sequencing analysis.

Jumpei Ito and Spyros Lytras performed phylogenetic analyses.

Jumpei Ito performed statistical, modelling, and bioinformatics analyses.

Jumpei Ito, Akatsuki Saito, Keita Matsuno, Kazuo Takayama, Shinya Tanaka, Takasuke Fukuhara, Terumasa Ikeda and Kei Sato designed the experiments and interpreted the results.

Jumpei Ito and Kei Sato wrote the original manuscript.

All authors reviewed and proofread the manuscript.

The Genotype to Phenotype Japan (G2P-Japan) Consortium contributed to the project administration.

## Acknowledgments

We would like to thank all members belonging to The Genotype to Phenotype Japan (G2P-Japan) Consortium. We thank Dr. Kenzo Tokunaga (National Institute for Infectious Diseases, Japan) and Dr. Jin Gohda (The University of Tokyo, Japan) for providing reagents. We also thank National Institute for Infectious Diseases, Japan for providing clinical isolates of BQ.1.1 (strain TY41-796-P1; GISAID ID: EPI_ISL_15579783) and BA.2 (strain TY40-385; GISAID ID: EPI_ISL_9595859). We appreciate the technical assistance from The Research Support Center, Research Center for Human Disease Modeling, Kyushu University Graduate School of Medical Sciences. We gratefully acknowledge all data contributors, i.e. the Authors and their Originating laboratories responsible for obtaining the specimens, and their Submitting laboratories for generating the genetic sequence and metadata and sharing via the GISAID Initiative, on which this research is based. The super-computing resource was provided by Human Genome Center at The University of Tokyo.

This study was supported in part by AMED SCARDA Japan Initiative for World-leading Vaccine Research and Development Centers “UTOPIA” (JP223fa627001, to Kei Sato), AMED SCARDA Program on R&D of new generation vaccine including new modality application (JP223fa727002, to Kei Sato); AMED Research Program on Emerging and Re-emerging Infectious Diseases (JP21fk0108574, to Hesham Nasser; JP21fk0108465, to Akatsuki Saito; JP21fk0108493, to Takasuke Fukuhara; JP22fk0108617 to Takasuke Fukuhara; JP22fk0108146, to Kei Sato; JP21fk0108494 to G2P-Japan Consortium, Keita Matsuno, Shinya Tanaka, Terumasa Ikeda, Takasuke Fukuhara, and Kei Sato; JP21fk0108425, to Kazuo Takayama and Kei Sato; JP21fk0108432, to Kazuo Takayama, Takasuke Fukuhara and Kei Sato); AMED Research Program on HIV/AIDS (JP22fk0410033, to Akatsuki Saito; JP22fk0410047, to Akatsuki Saito; JP22fk0410055, to Terumasa Ikeda; and JP22fk0410039, to Kei Sato); AMED CRDF Global Grant (JP22jk0210039 to Akatsuki Saito); AMED Japan Program for Infectious Diseases Research and Infrastructure (JP22wm0325009, to Akatsuki Saito; JP22wm0125008 to Keita Matsuno); AMED CREST (JP21gm1610005, to Kazuo Takayama; JP22gm1610008, to Takasuke Fukuhara); JST PRESTO (JPMJPR22R1, to Jumpei Ito); JST CREST (JPMJCR20H4, to Kei Sato); JSPS KAKENHI Grant-in-Aid for Scientific Research C (22K07103, to Terumasa Ikeda); JSPS KAKENHI Grant-in-Aid for Scientific Research B (21H02736, to Takasuke Fukuhara); JSPS KAKENHI Grant-in-Aid for Early-Career Scientists (22K16375, to Hesham Nasser; 20K15767, Jumpei Ito); JSPS Core-to-Core Program (A. Advanced Research Networks) (JPJSCCA20190008, to Kei Sato); JSPS Research Fellow DC2 (22J11578, to Keiya Uriu); JSPS Leading Initiative for Excellent Young Researchers (LEADER) (to Terumasa Ikeda); World-leading Innovative and Smart Education (WISE) Program 1801 from the Ministry of Education, Culture, Sports, Science and Technology (MEXT) (to Naganori Nao); The Tokyo Biochemical Research Foundation (to Kei Sato); Takeda Science Foundation (to Terumasa Ikeda); Mochida Memorial Foundation for Medical and Pharmaceutical Research (to Terumasa Ikeda); The Naito Foundation (to Terumasa Ikeda); Shin-Nihon Foundation of Advanced Medical Research (to Terumasa Ikeda); Waksman Foundation of Japan (to Terumasa Ikeda); an intramural grant from Kumamoto University COVID-19 Research Projects (AMABIE) (to Terumasa Ikeda); the UK’s Medical Research Council (to Spyros Lytras); and the project of National Institute of Virology and Bacteriology, Programme EXCELES, funded by the European Union, Next Generation EU (LX22NPO5103, to Jiri Zahradnik).

## Consortia

Saori Suzuki^2^, Marie Kato^8^, Zannatul Ferdous^8^, Hiromi Mouri^8^, Kenji Shishido^8^, Naoko Misawa^1^, Izumi Kimura^1^, Yusuke Kosugi^1^, Pan Lin^1^, Mai Suganami^1^, Mika Chiba^1^, Ryo Yoshimura^1^, Kyoko Yasuda^1^, Keiko Iida^1^, Naomi Ohsumi^1^, Adam P. Strange^1^, Daniel Sauter^1,30^, So Nakagawa^31^ Jiaqi Wu^31^, Yukio Watanabe^7^, Ayaka Sakamoto^7^, Naoko Yasuhara^7^, Takao Hashiguchi^32^, Tateki Suzuki^32^, Kanako Kimura^32^, Jiei Sasaki^32^, Yukari Nakajima^32^, Hisano Yajima^32^, Kotaro Shirakawa^32^, Akifumi Takaori-Kondo^32^, Kayoko Nagata^32^, Yasuhiro Kazuma^32^, Ryosuke Nomura^32^, Yoshihito Horisawa^32^, Yusuke Tashiro^32^, Yugo Kawa^32^, Takashi Irie^33^, Ryoko Kawabata^33^, Ryo Shimizu^12^, Otowa Takahashi^12^, Kimiko Ichihara^12^, Chihiro Motozono^34^, Mako Toyoda^34^, Takamasa Ueno^34^, Yuki Shibatani^14^, Tomoko Nishiuchi^14^

^30^University Hospital Tübingen, Tübingen, Germany

^31^Tokai University School of Medicine, Isehara, Japan

^32^Kyoto University, Kyoto, Japan

^33^Hiroshima University, Hiroshima, Japan

^34^Kumamoto University, Kumamoto, Japan

## Figure legends

**Supplementary Table 1.** Information on the number of estimated substitution events in each Omicron lineage

**Supplementary Table 2.** Effect size of each S substitution on R_e_ estimated by a hierarchal Bayesian model

**Supplementary Table 3.** Relative R_e_ value for a viral group represented by each S haplotype

**Supplementary Table 4.** Human sera used in this study

**Supplementary Table 5.** Primers used in this study

**Supplementary Table 6.** Summary of unexpected amino acid mutations detected in the working virus stocks

**Extended Data Fig. 1. Convergent evolution of the Omicron lineages.**

**a,** Detected substitution events at the convergent sites (related to **Fig. 1d,e**). Raw counts (left) and counts per 1 million (M) analyzed sequences (right) are shown. Unlike **Fig. 1d,e**, results for BA.2.75 are shown in addition to BA.1, BA.2, BA.4, and BA.5.

**b,** The co-occurrence network of S substitutions in the Omicron lineages. In the S haplotype dataset, a pair of substitutions with Pearson’s correlation > 0.9 is considered co-occurring substitutions and indicated as a link in the network. In the modeling analysis, a group of co-occurring substitutions was clustered, and one effect value was estimated for each substitution cluster.

**c,** Effect size of each S substitution on R_e_ (related to **Fig. 1f**). A dot and line indicate the posterior mean and the 95% Bayesian confidential interval (CI), respectively.

**d,** Relative R_e_ value for a viral group represented by each S haplotype, assuming a fixed generation time of 2.1 days (related to **Fig. 1g**). A dot and line indicate the posterior mean and the 95% Bayesian CI, respectively. Unlike **Fig. 1g**, the profile of all S substitutions analyzed is shown on the left side.

**Extended Data Fig. 2. Histological observations in infected hamsters**

**a,** IHC of the viral N protein in the middle portion of the tracheas of all infected hamsters at 2 d.p.i (4 hamsters per infection group). Each panel shows a representative result from an individual infected hamster.

**b,** IHC of the SARS-CoV-2 N protein in the lungs of infected hamsters at 2 d.p.i. (left) and 5 d.p.i (right) (4 hamsters per infection group). In each panel, IHC staining (top) and the digitalized N-positive area (bottom, indicated in red) are shown. The red numbers in the bottom panels indicate the percentage of the N-positive area. Summarized data are shown in **Fig.4c**. In **a and b**, N-positive cells are shown in brown. Scale bars, 1 mm (**a**); 5 mm (**b**).

## Methods

### Ethics statement

All experiments with hamsters were performed in accordance with the Science Council of Japan’s Guidelines for the Proper Conduct of Animal Experiments. The protocols were approved by the Institutional Animal Care and Use Committee of National University Corporation Hokkaido University (approval ID: 20-0123 and 20-0060). All protocols involving specimens from human subjects recruited at Interpark Kuramochi Clinic was reviewed and approved by the Institutional Review Board of Interpark Kuramochi Clinic (approval ID: G2021-004). All human subjects provided written informed consent. All protocols for the use of human specimens were reviewed and approved by the Institutional Review Boards of The Institute of Medical Science, The University of Tokyo (approval IDs: 2021-1-0416 and 2021-18-0617) and University of Miyazaki (approval ID: O-1021).

### Human serum collection

Convalescent sera were collected from fully vaccinated individuals who had been infected with BA.2 (9 2-dose vaccinated and 5 3-dose vaccinated; 11–61 days after testing. n=14 in total; average age: 47 years, range: 24–84 years, 64% male) (**Fig. 2a**), and fully vaccinated individuals who had been infected with BA.5 (2 2-dose vaccinated, 17 3-dose vaccinated and 1 4-dose vaccinated; 10–23 days after testing. n=20 in total; average age: 51 years, range: 25–73 years, 45% male) (**Fig. 2b**). The SARS-CoV-2 variants were identified as previously described^2,15,18^. Sera were inactivated at 56°C for 30 minutes and stored at –80°C until use. The details of the convalescent sera are summarized in **Supplementary Table 4**.

### Cell culture

HEK293T cells (a human embryonic kidney cell line; ATCC, CRL-3216), HEK293 cells (a human embryonic kidney cell line; ATCC, CRL-1573) and HOS-ACE2/TMPRSS2 cells (HOS cells stably expressing human ACE2 and TMPRSS2)^32,33^ were maintained in DMEM (high glucose) (Sigma-Aldrich, Cat# 6429-500ML) containing 10% fetal bovine serum (FBS, Sigma-Aldrich Cat# 172012-500ML) and 1% penicillin–streptomycin (PS) (Sigma-Aldrich, Cat# P4333-100ML). HEK293-ACE2 cells (HEK293 cells stably expressing human ACE2)^19^ were maintained in DMEM (high glucose) containing 10% FBS, 1 μg/ml puromycin (InvivoGen, Cat# ant-pr-1) and 1% PS. HEK293-ACE2/TMPRSS2 cells (HEK293 cells stably expressing human ACE2 and TMPRSS2)^19^ were maintained in DMEM (high glucose) containing 10% FBS, 1 μg/ml puromycin, 200 μg/ml hygromycin (Nacalai Tesque, Cat# 09287-84) and 1% PS. Vero cells [an African green monkey *(Chlorocebus sabaeus)* kidney cell line; JCRB Cell Bank, JCRB0111] were maintained in Eagle’s minimum essential medium (EMEM) (Sigma-Aldrich, Cat# M4655-500ML) containing 10% FBS and 1% PS. VeroE6/TMPRSS2 cells (VeroE6 cells stably expressing human TMPRSS2; JCRB Cell Bank, JCRB1819)^34^ were maintained in DMEM (low glucose) (Wako, Cat# 041-29775) containing 10% FBS, G418 (1 mg/ml; Nacalai Tesque, Cat# G8168-10ML) and 1% PS. Calu-3 cells (ATCC, HTB-55) were maintained in Eagle’s minimum essential medium (EMEM) (Sigma-Aldrich, Cat# M4655-500ML) containing 10% FBS and 1% PS. Calu-3/DSP_1-7_ cells (Calu-3 cells stably expressing DSP_1-7_)^35^ were maintained in EMEM (Wako, Cat# 056-08385) containing 20% FBS and 1% PS. Human airway and lung epithelial cells derived from human induced pluripotent stem cells (iPSCs) were manufactured according to established protocols as described below (see “Preparation of human airway and lung epithelial cells from human iPSCs” section) and provided by HiLung Inc. AO-ALI model was generated according to established protocols as described below (see “AO-ALI model” section).

### Viral genome sequencing

Viral genome sequencing was performed as previously described^15^. Briefly, the virus sequences were verified by viral RNA-sequencing analysis. Viral RNA was extracted using a QIAamp viral RNA mini kit (Qiagen, Cat# 52906). The sequencing library employed for total RNA sequencing was prepared using the NEBNext Ultra RNA Library Prep Kit for Illumina (New England Biolabs, Cat# E7530). Paired-end 76-bp sequencing was performed using a MiSeq system (Illumina) with MiSeq reagent kit v3 (Illumina, Cat# MS-102-3001). Sequencing reads were trimmed using fastp v0.21.0^36^ and subsequently mapped to the viral genome sequences of a lineage B isolate (strain Wuhan-Hu-1; GenBank accession number: NC_045512.2)^34^ using BWA-MEM v0.7.17^37^. Variant calling, filtering, and annotation were performed using SAMtools v1.9^38^ and snpEff v5.0e^39^.

### Phylogenetic reconstruction

A total of 5,345,749 SARS-CoV-2 genome sequences labelled as ‘Omicron’ and their corresponding metadata were retrieved from the GISAID database on October 3, 2022 (https://www.gisaid.org/)^40^. The dataset was then filtered based on the following criteria: i) only ‘original passage’ sequences, ii) collection date in 2022, iii) host labelled as ‘Human’, iv) sequence length above 28,000 base pairs and v) proportion of ambiguous bases below 2%. This filtering reduced the dataset to a total of 3,840,308 sequences. To ensure that PANGO lineage definitions in our dataset’s metadata included the latest circulating lineages, the GISAID metadata were downloaded again on October 15^th^, 2022, and PANGO lineages of our sequences were updated accordingly.

To construct an ML tree of Omicron lineages (**Fig. 1a**), we randomly sampled 100 sequences from BA.1, BA.2, BA.4, and BA.5 and 20 sequences from BA.2.75 and BQ.1.1. In addition, an outgroup sequence, EPI_ISL_466615, representing the oldest isolate of B.1.1 obtained in the UK was added to the dataset. The viral genome sequences were mapped to the reference sequence of Wuhan-Hu-1 (GenBank accession number: NC_045512.2) using Minimap2 v2.17^41^ and subsequently converted to a multiple sequence alignment according to the GISAID phylogenetic analysis pipeline

(https://github.com/roblanf/sarscov2phylo). The alignment sites corresponding to the 1–265 and 29674–29903 positions in the reference genome were masked (i.e., converted to NNN). Alignment sites at which >50% of sequences contained a gap or undetermined/ambiguous nucleotide were trimmed using trimAl v1.2^42^. Phylogenetic tree construction was performed via a three-step protocol: i) the first tree was constructed; ii) tips with longer external branches (Z score > 4) were removed from the dataset; iii) and the final tree was constructed. Tree reconstruction was performed by RAxML v8.2.12^43^ under the GTRCAT substitution model. The node support value was calculated by 100 times bootstrap analysis.

A separate phylogenetic tree was reconstructed for each Omicron lineage (BA.1, BA.2, BA.4 and BA.5) including all their descendant sublineages (**Fig. 1c**). To remove redundant sequences and reduce the volume of data for each reconstruction, a representative subsampling approach was used. 3,000 sequences from each Omicron lineage that had no substitutions at the convergent sites in S: 346, 444, 452, 460 and 486 for BA.1 and BA.2 or no substitutions in sites 346, 444 and 460 for BA.4 and BA.5 were randomly sampled from each dataset, weighting the sampling by the frequency of each PANGO lineage in the dataset. In this way we included a sample of background sequences with no ‘additional’ substitutions in the sites of interest with PANGO lineage frequencies representative of the full dataset. It was also ensured that the selected sequences had no ambiguous bases in the S gene (checked between sequence positions 21,000 to 26,000) to avoid ambiguous residues in the sites of interest. Recombinant PANGO lineages were excluded from the analysis.

After collecting the subsampled set of background sequences for each lineage, a maximum of 30 randomly selected sequences of each PANGO sublineage with at least one additional substitution at the convergent sites were added to the dataset. This subsampling approach aimed to capture sequences of all sublineages that have acquired additional mutations at the convergent sites, while maintaining a large set of background lineages that reflects circulating lineage distribution. One SARS-CoV-2 sequence from the sister lineage of each set with a recent collection date was also added to each dataset to be used as an outgroup of the phylogeny [for the BA.1 tree, EPI_ISL_15170885 (BA.2); for the BA.2 tree, EPI_ISL_15148193 (BA.1); for the BA.4 tree, EPI_ISL_15192101 (BA.5); and for BA.5 the tree, EPI_ISL_15174939 (BA.4)].

Each lineage sequence dataset was aligned using the ‘global_profile_alignment.sh’ from the SARS-CoV-2 global phylogeny pipeline [www.doi.org/10.5281/zenodo.3958883], utilising MAFFT^44^. Phylogenies were reconstructed using iqtree2 (v2.1.3)^45^ under a GTR+I+F+G4 model with 1000 ultrafast bootstrap replicates to determine node support. Trees were manually rerooted on the branch leading to the outgroup sequence and time-calibrated with TreeTime^46^ (with the ‘--keep-root’ option to preserve the outgroup rooting). Branches leading to tips with dates not matching the root-to-tip regression model were removed from the phylogeny using the ete3 python package^47^. The final trees for BA.1, BA.2, BA.4 and BA.5 contain 3,901, 5,343, 3,328, and 5,197 sequences, respectively.

### Ancestral node reconstruction of site substitutions

To infer the branches where substitution events occurred at the five convergent sites (positioned at 346, 444, 452, 460, 486) in the trees of Omicron lineages, we reconstructed the ancestral state of the substitution profile at the convergent sites in each node using a parsimony method, implemented by the phangorn package (https://github.com/KlausVigo/phangorn). Internal nodes with substitution probabilities above or equal to 0.5 were annotated as having the substitution. Branches where substitutions took place for each site were denoted as branches connecting an ancestral internal node with no substitution to an internal node that has a substitution. Additionally, 70% of tips descending from that internal node were also required to have the substitution and at least 3 tips needed to be descended from the node, to avoid picking up branches with low support or clades that reverted back to the original residue. The analysis was performed on R v4.1.2 (https://www.r-project.org/).

### Modeling the relationship between viral epidemic dynamics and S substitutions

Motivated by the model established by Obermeyer et al^48^, we developed a method to model the relationship between viral epidemic dynamics and S substitutions. This model can simultaneously estimate i) the effect of each S substitution on R_e_ and ii) the relative R_e_ of a viral group represented by each S haplotype. The key concept of the model used in this study is the same as the one in Obermeyer et al^48^. However, our method is independent of the predefined viral classification such as PANGO lineage but based on the viral classification according to the profile of S substitutions. Therefore, our method can link the effect of S substitutions to viral epidemic dynamics in a more direct manner. Also, in our method, a Markov Chain Monte Carlo (MCMC) method is used for parameter estimation instead of variational inference, an approximation method.

The data used in this analysis were downloaded from the GISAID database (https://www.gisaid.org/) on November 7, 2022. For quality control, we excluded the data of viral sequences with the following features from the analysis: i) a lack of collection date information; ii) sampling in animals other than humans; iii) >1% undetermined nucleotide characters; or iv) sampling by quarantine. Furthermore, in this analysis, we analyzed viral sequences of the Omicron lineages collected in the UK from March 1, 2022, to October 15^th^, 2022.

We selected S substitutions (including insertions and deletions) to be analyzed and classified Omicron sequences into S haplotypes according to the profile of the selected S substitutions: We analyzed S substitutions observed in ≥200 sequences in the dataset we used. We excluded S substitutions commonly (≥90%) detected in sequences analyzed. According to the criteria above, 123 S substitutions were retrieved. Subsequently, we classified the sequences according to the profile of S substitutions above (referred to as S haplotype). We excluded S haplotypes with ≤30 sequences from the downstream analyses. According to the criterion above, 254 S haplotypes, composed of 375,121 sequences, were retrieved. The substitution profile was represented as a matrix, where the rows and columns depict S haplotypes and S substitutions, respectively. An element in the matrix represents the status [presence (1) or absence (0)] of one S substitution in one S haplotype. Next, we identified a group of highly co-occurring substitutions (i.e., a pair of substitutions with >0.9 Pearson’s correlation in the substitution profile matrix) and clustered these substitutions as a substitution cluster (**Extended Data Fig. 1b**). For example, the L452R: F486V cluster represents the L452R and F486V substitutions. For one substitution cluster, the mean value of the substitution statuses (0 or 1) of the members of substitutions was calculated for each S haplotype, and the mean value was used as the substitution status of the substitution cluster. For example, if one S haplotype has L452R but not F486V, the substitution status of the L452R:F486V cluster of the haplotype was set at 0.5. Consequently, our dataset included the profile of 107 S substitutions/substitution clusters for 254 S haplotypes. Next, to set the major S haplotype of BA.2 as the reference S haplotype (or lineage) in the statistical model described below, we transformed the S substitution profile matrix by subtracting the substitution profile of the major S haplotype of BA.2 from those for all S haplotypes. Consequently, elements in the transformed S substitution profile matrix were converted to −1, 0, or 1: The zero value means that the status of a substitution in one haplotype is the same as that in the reference haplotype. The one value means that a substitution is present in one haplotype but not in the reference haplotype. The minus one value means that a substitution is absent in one haplotype but present in the reference haplotype. As a consequence of the transformation, the relative R_e_ value for the reference haplotype was set at 1 in the parameter estimation in the statistical model described below. Finally, the number of viral sequences belonging to each S haplotype collected on each day was counted, and the count matrix was constructed as an input for the statistical model described below.

We assigned one major lineage classification (i.e., BA.1 BA.2, BA.4, BA.5, and BA.2.75) to each S haplotype: We examined the major lineage classification of respective viral sequences belonging to one S haplotype, and the classification of the S haplotype was determined according to the majority vote system.

We constructed a Bayesian hierarchal model, which represents the epidemic dynamics of each S haplotype according to growth rate parameters for each S haplotype, which is represented by a linear combination of the effect of S substitutions. Arrays in the model index over one or more indices: L = 254 viral lineages (i.e., S haplotypes) *l*; S = 107 substitutions/substitution clusters *s*; and T = 229 days *t*. The model is:

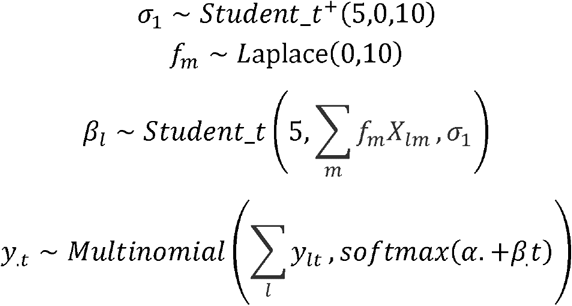

The count of viral lineage *l* at time *t, y_lt_*, is modeled as a hierarchal Multinomial logistic regression with intercept *α_l_* and slope *β_l_* parameters for lineage *l*. The slope (or viral lineage growth) parameter *β_l_* is generated from Student’s t distribution with five degrees of freedom, the mean value represented by *f_m_X_lm_*, and standard deviation, *σ*_1_. *f_m_X_lm_* denotes the linear combination of the effect of each substitution, where *f_m_* and *X_lm_* are the effect of substitution *m* and the profile of substitution *m* in lineage *l* (i.e., the substitution profile matrix constructed in the above paragraph), respectively. As a prior of *f_m_*, the Laplace distribution with the mean 0 and the standard deviation 10 was set. In other words, we estimated the parameter *f_m_* in the framework of Bayesian least absolute shrinkage and selection operator (LASSO). As a prior of *σ*_1_, a half Student’s t distribution with the mean 0 and the standard deviation 10 was set. For the other parameters, non-informative priors were set.

The relative R_e_ of each viral lineage, *r_l_*, was calculated according to the slope parameter *β_l_* as:

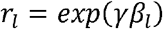

where *γ* is the average viral generation time (2.1 days) (http://sonorouschocolate.com/covid19/index.php?title=Estimating_Generation_Time_Of_Omicron). Similarly, the effect size of substitution *m* on the relative R_e_, *F_l_*, was calculated according to the coefficient as:

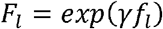

Parameter estimation was performed via the MCMC approach implemented in CmdStan v2.30.1 (https://mc-stan.org) with CmdStanr v0.5.3 (https://mc-stan.org/cmdstanr/). Four independent MCMC chains were run with 500 and 2,000 steps in the warmup and sampling iterations, respectively. We confirmed that all estimated parameters showed <1.01 R-hat convergence diagnostic values and >200 effective sampling size values, indicating that the MCMC runs were successfully convergent. The above analyses were performed in R v4.2.1 (https://www.r-project.org/). Information on the estimated effect size of each substitution or substitution cluster on relative R_e_ and relative R_e_ for each S haplotype are summarized in **Supplementary Table 2,3**.

Since our model simply represents the viral lineage growth parameter (*β_l_*) as the linear combination of the effects of S substitutions, the model can predict the total effect of a set of substations on relative R_e_. Using this property of the model, we predicted the total effect of substitutions at the convergent sites (**Fig. 1h**) and the ancestral relative viral fitness for each node in the BA.5 tree (**Fig. 1i**). We calculated these values as the sum of the posterior means of the effects of substitutions of interest. To reconstruct the ancestral relative viral fitness of each node of the BA.5 tree, we first reconstructed the ancestral state of the S substitution profile in each node of the tree using a parsimony method, implemented by the phangorn package. Subsequently, we predicted the relative viral fitness for each node according to the reconstructed ancestral mutation profile for the node. The above analyses were performed in R v4.2.1 (https://www.r-project.org/).

### Plasmid construction

Plasmids expressing the codon-optimized SARS-CoV-2 S proteins of B.1.1 (the parental D614G-bearing variant), BA.2 and BA.5, and BA.2.75 were prepared in our previous studies^27^. Plasmids expressing the codon-optimized S proteins of BQ.1.1, BA.5 S-based derivatives and BA.2 S-based derivatives were generated by site-directed overlap extension PCR using the primers listed in **Supplementary Table 5**. The resulting PCR fragment was digested with KpnI (New England Biolabs, Cat# R0142S) and NotI (New England Biolabs, Cat# R1089S) and inserted into the corresponding site of the pCAGGS vector^49^.

Nucleotide sequences were determined by DNA sequencing services (Eurofins), and the sequence data were analyzed by Sequencher v5.1 software (Gene Codes Corporation). Plasmids for yeast surface display were constructed by restriction enzyme-free cloning by incorporation of RBD genes [“construct 3” in ref.^23^, covering residues 330–528] into the pJYDC1 plasmid (Addgene, Cat# 162458). The primers are listed in **Supplementary Table 6**. The non-mutated RBD genes (BA.2, BA.5, and BQ.1) were purchased from Twist Biosciences.

### Neutralization assay

Pseudoviruses were prepared as previously described^2,15,18,19,24,33,50–52^. Briefly, lentivirus (HIV-1)-based, luciferase-expressing reporter viruses were pseudotyped with SARS-CoV-2 S proteins. HEK293T cells (1,000,000 cells) were cotransfected with 1 μg psPAX2-IN/HiBiT^32^, 1 μg pWPI-Luc2^32^, and 500 ng plasmids expressing parental S or its derivatives using PEI Max (Polysciences, Cat# 24765-1) according to the manufacturer’s protocol. Two days posttransfection, the culture supernatants were harvested and centrifuged. The pseudoviruses were stored at −80°C until use.

The neutralization assay (**Fig. 2**) was prepared as previously described^2,15,18,24,33,50–52^. Briefly, the SARS-CoV-2 S pseudoviruses (counting ~20,000 relative light units) were incubated with serially diluted (120-fold to 87,480-fold dilution at the final concentration) heat-inactivated sera at 37°C for 1 hour. Pseudoviruses without sera were included as controls. Then, a 40 μl mixture of pseudovirus and serum/antibody was added to HOS-ACE2/TMPRSS2 cells (10,000 cells/50 μl) in a 96-well white plate. At 2 d.p.i., the infected cells were lysed with a One-Glo luciferase assay system (Promega, Cat# E6130), a Bright-Glo luciferase assay system (Promega, Cat# E2650), or a britelite plus Reporter Gene Assay System (PerkinElmer, Cat# 6066769), and the luminescent signal was measured using a GloMax explorer multimode microplate reader 3500 (Promega) or CentroXS3 (Berthhold Technologies). The assay of each serum sample was performed in triplicate, and the 50% neutralization titer (NT_50_) was calculated using Prism 9 software v9.1.1 (GraphPad Software).

### SARS-CoV-2 preparation and titration

The working virus stocks of SARS-CoV-2 were prepared and titrated as previously described^15,18,19,53^. In this study, clinical isolates of B.1.1 (strain TKYE610670; GISAID ID: EPI_ISL_479681)^17^, Delta (B.1.617.2, strain TKYTK1734; GISAID ID: EPI_ISL_2378732)^16^, BA.2 (strain TY40-385; GISAID ID: EPI_ISL_9595859)^15^ and BA.5 (strain TKYS14631; GISAID ID: EPI_ISL_12812500)^2,54^, and BQ.1.1 (strain TY41-796-P1; GISAID ID: EPI_ISL_15579783) were used. In brief, 20 μl of the seed virus was inoculated into VeroE6/TMPRSS2 cells (5,000,000 cells in a T-75 flask). One h.p.i., the culture medium was replaced with DMEM (low glucose) (Wako, Cat# 041-29775) containing 2% FBS and 1% PS. At 3 d.p.i., the culture medium was harvested and centrifuged, and the supernatants were collected as the working virus stock.

The titer of the prepared working virus was measured as the 50% tissue culture infectious dose (TCID50). Briefly, one day before infection, VeroE6/TMPRSS2 cells (10,000 cells) were seeded into a 96-well plate. Serially diluted virus stocks were inoculated into the cells and incubated at 37°C for 4 days. The cells were observed under a microscope to judge the CPE appearance. The value of TCID_50_/ml was calculated with the Reed–Muench method^55^.

For verification of the sequences of SARS-CoV-2 working viruses, viral RNA was extracted from the working viruses using a QIAamp viral RNA mini kit (Qiagen, Cat# 52906) and viral genome sequences were analyzed as described above (see “Viral genome sequencing” section). Information on the unexpected substitutions detected is summarized in **Supplementary Table 6**, and the raw data are deposited in the GitHub repository (https://github.com/TheSatoLab/BQ.1).

### Yeast surface display

Yeast surface display (**Fig. 3a**) was performed as previously described^2,22,23^. Briefly, the *S. cerevisiae* EBY100 yeasts were transformed with RBD expression plasmid and grown (220 rpm, 30°C, SD-CAA media). The expression media 1/9 (ref.^56^) was inoculated to starting OD_600_ 0.7–1 by overnight grown culture and cultivated for 24 hours at 20°C. The expression media was supplemented with 10 nM DMSO solubilized bilirubin (Sigma-Aldrich, Cat# 14370-1G) for activation of eUnaG2 fluorescence (excitation at 498 nm, emission at 527 nm).

Yeast cells were washed in ice-cold PBSB buffer (PBS with 1 mg/ml BSA), liquated (100 μl), transferred in an analysis solution and incubated for 8 hours. The analysis solutions consisted of a series of CF®640R succinimidyl ester labeled (Biotium, Cat# 92108) ACE2 peptidase domain (residues 18–740) concentrations, PBSB buffer and 1 nM bilirubin. Incubated samples were washed twice with PBSB buffer and transferred into a 96-well plate (Thermo Fisher Scientific, Cat# 268200) for automated data acquisition by a CytoFLEX S Flow Cytometer (Beckman Coulter, USA, Cat#. N0-V4-B2-Y4). The gating and analysis strategies were described previously^23^. The titration curves were fitted with nonlinear least-squares regression using Python v3.7 and two additional parameters to describe the titration curve^23^.

### Pseudovirus infection

Pseudovirus infection (**Fig. 3b**) was performed as previously described^2,15,18,19,24,33,50–52^. Briefly, the amount of pseudoviruses prepared was quantified by the HiBiT assay using a Nano Glo HiBiT lytic detection system (Promega, Cat# N3040) as previously described^32,57^. For measurement of pseudovirus infectivity, the same amount of pseudoviruses (normalized to the HiBiT value, which indicates the amount of HIV-1 p24 antigen) was inoculated into HOS-ACE2/TMPRSS2 cells, HEK293-ACE2 cells or HEK293-ACE2/TMPRSS2 cells and viral infectivity was measured as described above (see “Neutralization assay” section). For analysis of the effect of TMPRSS2 on pseudovirus infectivity (**Fig. 3c**), the fold change of the values of HEK293-ACE2/TMPRSS2 to HEK293-ACE2 was calculated.

### SARS-CoV-2 S-based fusion assay

A SARS-CoV-2 S-based fusion assay (**Fig. 3d–f**) was performed as previously described^2,15–19,25,53^. Briefly, on day 1, effector cells (i.e., S-expressing cells) and target cells (Calu-3/DSP_1-7_ cells) were prepared at a density of 0.6–0.8 × 10^6^ cells in a 6-well plate. On day 2, for the preparation of effector cells, HEK293 cells were cotransfected with the S expression plasmids (400 ng) and pDSP_8-11_ (ref.^58^) (400 ng) using TransIT-LT1 (Takara, Cat# MIR2300). On day 3 (24 hours posttransfection), 16,000 effector cells were detached and reseeded into a 96-well black plate (PerkinElmer, Cat# 6005225), and target cells were reseeded at a density of 1,000,000 cells/2 ml/well in 6-well plates. On day 4 (48 hours posttransfection), target cells were incubated with EnduRen live cell substrate (Promega, Cat# E6481) for 3 hours and then detached, and 32,000 target cells were added to a 96-well plate with effector cells. *Renilla* luciferase activity was measured at the indicated time points using Centro XS3 LB960 (Berthhold Technologies). For measurement of the surface expression level of the S protein, effector cells were stained with rabbit anti-SARS-CoV-2 S S1/S2 polyclonal antibody (Thermo Fisher Scientific, Cat# PA5-112048, 1:100). Normal rabbit IgG (Southern Biotech, Cat# 0111-01, 1:100) was used as a negative control, and APC-conjugated goat anti-rabbit IgG polyclonal antibody (Jackson ImmunoResearch, Cat# 111-136-144, 1:50) was used as a secondary antibody. The surface expression level of S proteins (**Fig. 3d**) was measured using a FACS Canto II (BD Biosciences) and the data were analyzed using FlowJo software v10.7.1 (BD Biosciences). For calculation of fusion activity, *Renilla* luciferase activity was normalized to the MFI of surface S proteins. The normalized value (i.e., *Renilla* luciferase activity per the surface S MFI) is shown as fusion activity.

### AO-ALI model

An airway organoid (AO) model was generated according to our previous report^2,59^. Briefly, normal human bronchial epithelial cells (NHBEs, Cat# CC-2540, Lonza) were used to generate AOs. NHBEs were suspended in 10 mg/ml cold Matrigel growth factor reduced basement membrane matrix (Corning, Cat# 354230). Fifty microliters of cell suspension were solidified on prewarmed cell culture-treated multiple dishes (24-well plates; Thermo Fisher Scientific, Cat# 142475) at 37°C for 10 min, and then, 500 μl of expansion medium was added to each well. AOs were cultured with AO expansion medium for 10 days. For maturation of the AOs, expanded AOs were cultured with AO differentiation medium for 5 days.

The AO-ALI model (**Fig. 3j**) was generated according to our previous report^2,59^. For generation of AO-ALI, expanding AOs were dissociated into single cells, and then were seeded into Transwell inserts (Corning, Cat# 3413) in a 24-well plate. AO-ALI was cultured with AO differentiation medium for 5 days to promote their maturation. AO-ALI was infected with SARS-CoV-2 from the apical side.

### Preparation of human airway and alveolar epithelial cells from human iPSCs

The air-liquid interface culture of airway and alveolar epithelial cells (**Fig. 3k,l**) was differentiated from human iPSC-derived lung progenitor cells as previously described^2,15,54,60–62^. Briefly, alveolar progenitor cells were induced stepwise from human iPSCs according to a 21-day and 4-step protocol^60^. At day 21, alveolar progenitor cells were isolated with the specific surface antigen carboxypeptidase M and seeded onto the upper chamber of a 24-well Cell Culture Insert (Falcon, #353104), followed by 28-day and 7-day differentiation of airway and alveolar epithelial cells, respectively. Alveolar differentiation medium with dexamethasone (Sigma-Aldrich, Cat# D4902), KGF (PeproTech, Cat# 100-19), 8-Br-cAMP (Biolog, Cat# B007), 3-isobutyl 1-methylxanthine (IBMX) (Fujifilm Wako, Cat# 095-03413), CHIR99021 (Axon Medchem, Cat# 1386), and SB431542 (Fujifilm Wako, Cat# 198-16543) was used for the induction of alveolar epithelial cells. PneumaCult ALI (STEMCELL Technologies, Cat# ST-05001) with heparin (Nacalai Tesque, Cat# 17513-96) and Y-27632 (LC Laboratories, Cat# Y-5301) hydrocortisone (Sigma-Aldrich, Cat# H0135) was used for induction of airway epithelial cells.

### Airway-on-a-chi ps

Airway-on-a-chips (**Fig. 3m**) were prepared as previously described^2,26,54^. Human lung microvascular endothelial cells (HMVEC-L) were obtained from Lonza (Cat# CC-2527) and cultured with EGM-2-MV medium (Lonza, Cat# CC-3202). For preparation of the airway-on-a-chip, first, the bottom channel of a polydimethylsiloxane (PDMS) device was precoated with fibronectin (3 μg/ml, Sigma-Aldrich, Cat# F1141). The microfluidic device was generated according to our previous report^63^. HMVEC-L cells were suspended at 5,000,000 cells/ml in EGM2-MV medium. Then, 10 μl of suspension medium was injected into the fibronectin-coated bottom channel of the PDMS device. Then, the PDMS device was turned upside down and incubated. After 1 hour, the device was turned over, and the EGM2-MV medium was added into the bottom channel. After 4 days, AOs were dissociated and seeded into the top channel. AOs were generated according to our previous report^59^. AOs were dissociated into single cells and then suspended at 5,000,000 cells/ml in the AO differentiation medium. Ten microliter suspension medium was injected into the top channel. After 1 hour, the AO differentiation medium was added to the top channel. In the infection experiments (**Fig. 3m**), the AO differentiation medium containing either BA.2, BA.5, BQ.1.1 or Delta isolate (500 TCID50) was inoculated into the top channel. At 2 h.p.i., the top and bottom channels were washed and cultured with AO differentiation and EGM2-MV medium, respectively. The culture supernatants were collected, and viral RNA was quantified using RT–qPCR (see “RT–qPCR” section above).

### Microfluidic device

A microfluidic device was generated according to our previous report^2,63^. Briefly, the microfluidic device consisted of two layers of microchannels separated by a semipermeable membrane. The microchannel layers were fabricated from PDMS using a soft lithographic method. PDMS prepolymer (Dow Corning, Cat# SYLGARD 184) at a base to curing agent ratio of 10:1 was cast against a mold composed of SU-8 2150 (MicroChem, Cat# SU-8 2150) patterns formed on a silicon wafer. The cross-sectional size of the microchannels was 1 mm in width and 330 μm in height. Access holes were punched through the PDMS using a 6-mm biopsy punch (Kai Corporation, Cat# BP-L60K) to introduce solutions into the microchannels. Two PDMS layers were bonded to a PET membrane containing 3.0-μm pores (Falcon, Cat# 353091) using a thin layer of liquid PDMS prepolymer as the mortar. PDMS prepolymer was spin-coated (4000 rpm for 60 sec) onto a glass slide. Subsequently, both the top and bottom channel layers were placed on the glass slide to transfer the thin layer of PDMS prepolymer onto the embossed PDMS surfaces. The membrane was then placed onto the bottom layer and sandwiched with the top layer. The combined layers were left at room temperature for 1 day to remove air bubbles and then placed in an oven at 60°C overnight to cure the PDMS glue. The PDMS devices were sterilized by placing them under UV light for 1 hour before the cell culture.

### SARS-CoV-2 infection

One day before infection, Vero cells (10,000 cells), VeroE6/TMPRSS2 cells (10,000 cells) and Calu-3 cells (10,000 cells) were seeded into a 96-well plate. SARS-CoV-2 [1,000 TCID50 for Vero cells (**Fig. 3g**); 100 TCID50 for VeroE6/TMPRSS2 cells (**Fig. 3h**) and Calu-3 cells (**Fig. 3i**)] was inoculated and incubated at 37°C for 1 hour. The infected cells were washed, and 180 μl of culture medium was added. The culture supernatant (10 μl) was harvested at the indicated timepoints and used for RT–qPCR to quantify the viral RNA copy number (see “RT–qPCR” section below). In the infection experiments using human iPSC-derived airway and lung epithelial cells (**Fig. 3k,l**), working viruses were diluted with Opti-MEM (Thermo Fisher Scientific, Cat# 11058021). The diluted viruses (1,000 TCID_50_ in 100□μl) were inoculated onto the apical side of the culture and incubated at 37□°C for 1 hour. The inoculated viruses were removed and washed twice with Opti-MEM. For collection of the viruses, 100□μl Opti-MEM was applied onto the apical side of the culture and incubated at 37□°C for 10□minutes. The Opti-MEM was collected and used for RT–qPCR to quantify the viral RNA copy number (see “RT–qPCR” section below). The infection experiments using an airway-on-a-chip system (**Fig. 3m**) were performed as described above (see “Airway-on-a-chips” section).

### RT–qPCR

RT–qPCR was performed as previously described^2,15–19,53,54,64^. Briefly, 5 μl culture supernatant was mixed with 5 μl of 2 × RNA lysis buffer [2% Triton X-100 (Nacalai Tesque, Cat# 35501-15), 50 mM KCl, 100 mM Tris-HCl (pH 7.4), 40% glycerol, 0.8 U/μl recombinant RNase inhibitor (Takara, Cat# 2313B)] and incubated at room temperature for 10 min. RNase-free water (90 μl) was added, and the diluted sample (2.5 μl) was used as the template for real-time RT-PCR performed according to the manufacturer’s protocol using One Step TB Green PrimeScript PLUS RT-PCR kit (Takara, Cat# RR096A) and the following primers: Forward *N*, 5’-AGC CTC TTC TCG TTC CTC ATC AC-3’; and Reverse *N*, 5’-CCG CCA TTG CCA GCC ATT C-3’. The viral RNA copy number was standardized with a SARS-CoV-2 direct detection RT-qPCR kit (Takara, Cat# RC300A). Fluorescent signals were acquired using a QuantStudio 1 Real-Time PCR system (Thermo Fisher Scientific), QuantStudio 3 Real-Time PCR system (Thermo Fisher Scientific), QuantStudio 5 Real-Time PCR system (Thermo Fisher Scientific), StepOne Plus Real-Time PCR system (Thermo Fisher Scientific), CFX Connect Real-Time PCR Detection system (Bio-Rad), Eco Real-Time PCR System (Illumina), qTOWER3 G Real-Time System (Analytik Jena) Thermal Cycler Dice Real Time System III (Takara) or 7500 Real-Time PCR System (Thermo Fisher Scientific).

### Animal experiments

Animal experiments (**Fig. 4 and Extended Data Fig. 2**) were performed as previously described^27^. Syrian hamsters (male, 4 weeks old) were purchased from Japan SLC Inc. (Shizuoka, Japan). For the virus infection experiments, hamsters were anesthetized by intramuscular injection of a mixture of 0.15 mg/kg medetomidine hydrochloride (Domitor®, Nippon Zenyaku Kogyo), 2.0 mg/kg midazolam (Dormicum®, Fujifilm Wako, Cat# 135-13791) and 2.5 mg/kg butorphanol (Vetorphale®, Meiji Seika Pharma) or 0.15 mg/kg medetomidine hydrochloride, 4.0 mg/kg alphaxaone (Alfaxan®, Jurox) and 2.5 mg/kg butorphanol. BA.5, BQ1.1 and Delta (10,000 TCID50 in 100 μl) or saline (100 μl) was intranasally inoculated under anesthesia. Oral swabs were collected at the indicated timepoints. Body weight was recorded daily by 7 d.p.i. Enhanced pause (Penh), the ratio of time to peak expiratory follow relative to the total expiratory time (Rpef) were measured every day until 7 d.p.i. (see below). Lung tissues were anatomically collected at 2 and 5 d.p.i. The viral RNA load in the oral swabs and respiratory tissues was determined by RT–qPCR. These tissues were also used for IHC and histopathological analyses (see below).

### Lung function test

Lung function tests (**Fig. 4a**) were routinely performed as previously described^2,15,17,18,54^. The two respiratory parameters (Penh and Rpef) were measured by using a Buxco Small Animal Whole Body Plethysmography system (DSI) according to the manufacturer’s instructions. In brief, a hamster was placed in an unrestrained plethysmography chamber and allowed to acclimatize for 30 seconds. Then, data were acquired over a 2.5-minute period by using FinePointe Station and Review software v2.9.2.12849 (DSI).

### Immunohistochemistry

Immunohistochemistry (IHC) (**Fig. 4c and Extended Data Fig. 2**) was performed as previously described^2,15,17,18,54^ using an Autostainer Link 48 (Dako). The deparaffinized sections were exposed to EnVision FLEX target retrieval solution high pH (Agilent, Cat# K8004) for 20 minutes at 97°C for activation, and a mouse anti-SARS-CoV-2 N monoclonal antibody (clone 1035111, R&D Systems, Cat# MAB10474-SP, 1:400) was used as a primary antibody. The sections were sensitized using EnVision FLEX for 15 minutes and visualized by peroxidase-based enzymatic reaction with 3,3’-diaminobenzidine tetrahydrochloride (Dako, Cat# DM827) as substrate for 5 minutes. The N protein positivity was evaluated by certificated pathologists as previously described^2,15,17,18,54^. Images were incorporated as virtual slides by NDP.scan software v3.2.4 (Hamamatsu Photonics). The N-protein positivity was measured as the area using Fiji software v2.2.0 (ImageJ).

### H&E staining

H&E staining (**Fig. 4d**) was performed as previously described^2,15,17,18,54^. Briefly, excised animal tissues were fixed with 10% formalin neutral buffer solution and processed for paraffin embedding. The paraffin blocks were sectioned at a thickness of 3 μm and then mounted on MAS-GP-coated glass slides (Matsunami Glass, Cat# S9901). H&E staining was performed according to a standard protocol.

### Histopathological scoring

Histopathological scoring (**Fig. 4e**) was performed as previously described^2,15,17,18,54^. Pathological features, including (i) bronchitis or bronchiolitis, (ii) hemorrhage with congestive edema, (iii) alveolar damage with epithelial apoptosis and macrophage infiltration, (iv) hyperplasia of type II pneumocytes, and (v) the area of hyperplasia of large type II pneumocytes, were evaluated by certified pathologists, and the degree of these pathological findings was arbitrarily scored using a four-tiered system as 0 (negative), 1 (weak), 2 (moderate), and 3 (severe). The “large type II pneumocytes” are type II pneumocytes with hyperplasia exhibiting more than 10-μm-diameter nuclei. We described “large type II pneumocytes” as one of the notable histopathological features of SARS-CoV-2 infection in our previous studies^2,15,17,18,54^. The total histological score is the sum of these five indices.

### Statistics and reproducibility

Statistical significance was tested using a two-sided Mann–Whitney *U* test, a two-sided Student’s *t* test, a two-sided Welch’s *t* test, or a two-sided paired *t-*test unless otherwise noted. The tests above were performed using Prism 9 software v9.1.1 (GraphPad Software).

In the time-course experiments (**Fig. 3e–m, 4a–b,e**), a multiple regression analysis including experimental conditions (i.e., the types of infected viruses) as explanatory variables and timepoints as qualitative control variables was performed to evaluate the difference between experimental conditions thorough all timepoints. The initial time point was removed from the analysis. The *P* value was calculated by a two-sided Wald test. Subsequently, familywise error rates (FWERs) were calculated by the Holm method. These analyses were performed on R v4.1.2 (https://www.r-project.org/).

Principal component analysis to representing the antigenicity of the S proteins was performed (**Fig. 2d**). The NT50 values for biological replicates were scaled, and subsequently, principal component analysis was performed using the prcomp function on R v4.1.2 (https://www.r-project.org/).

In **Fig. 4c,d, and Extended Data Fig. 2**, photographs shown are the representative areas of at least two independent experiments by using four hamsters at each timepoint.

## Data availability

All databases/datasets used in this study are available from the GISAID database (https://www.gisaid.org; EPI_SET_221203cz, EPI_SET_221203ep, EPI_SET_221203qr, EPI_SET_221203se, and EPI_SET_221203vk) and GenBank database (https://www.ncbi.nlm.nih.gov/genbank/). Viral genome sequencing data for working viral stocks are available in the GitHub repository (https://github.com/TheSatoLab/BQ.1).

## Code availability

The computational codes used in the present study and the GISAID supplemental tables for EPI_SET_221203cz, EPI_SET_221203ep, EPI_SET_221203qr, EPI_SET_221203se, and EPI_SET_221203vk are available in the GitHub repository (https://github.com/TheSatoLab/BQ.1).

