## Supplementary figures and images for "Convergent evolution of the SARS-CoV-2 Omicron subvariants leading to the emergence of BQ.1.1 variant"

### Extended Data Fig. 1

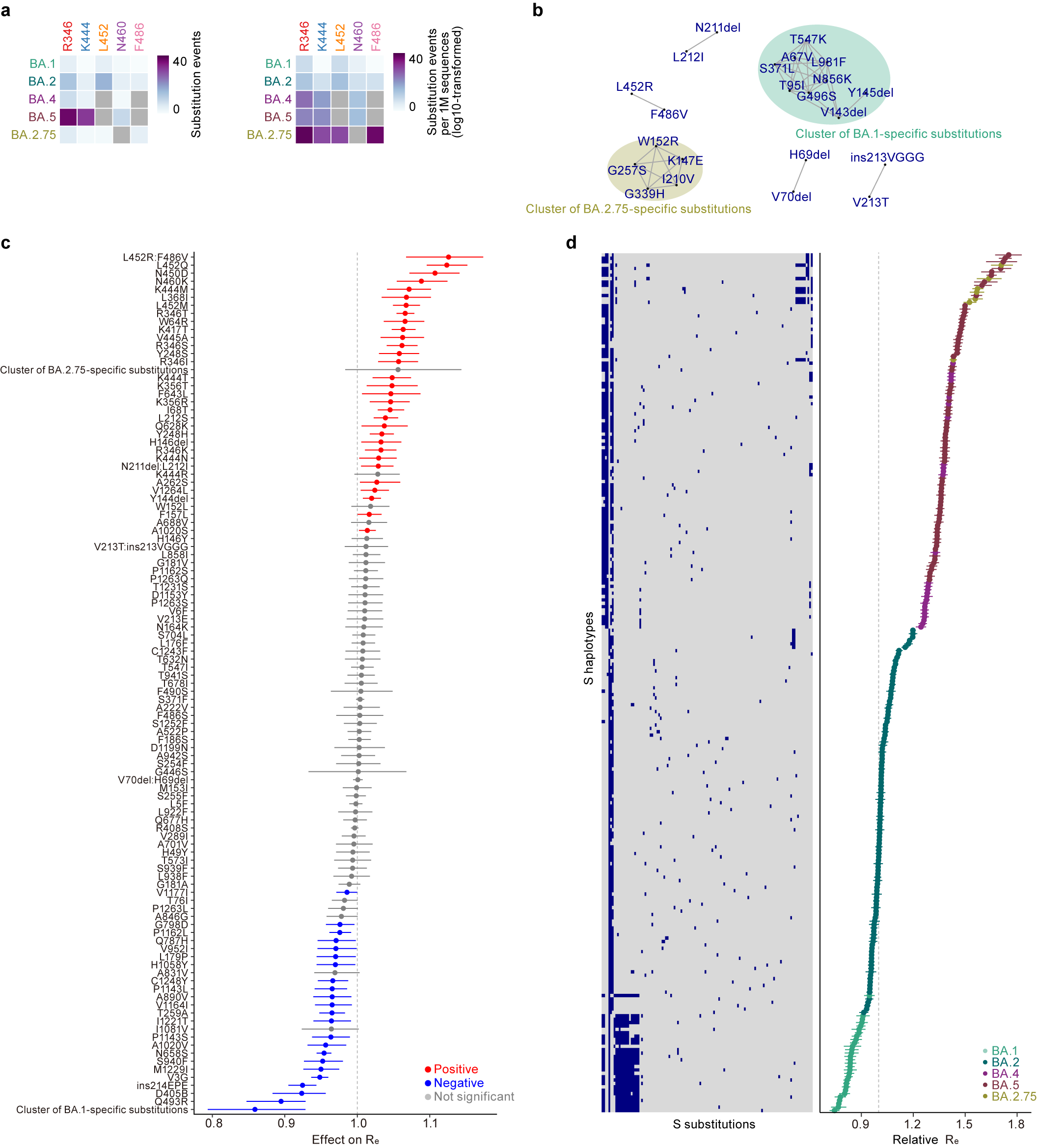

### Extended Data Fig. 2

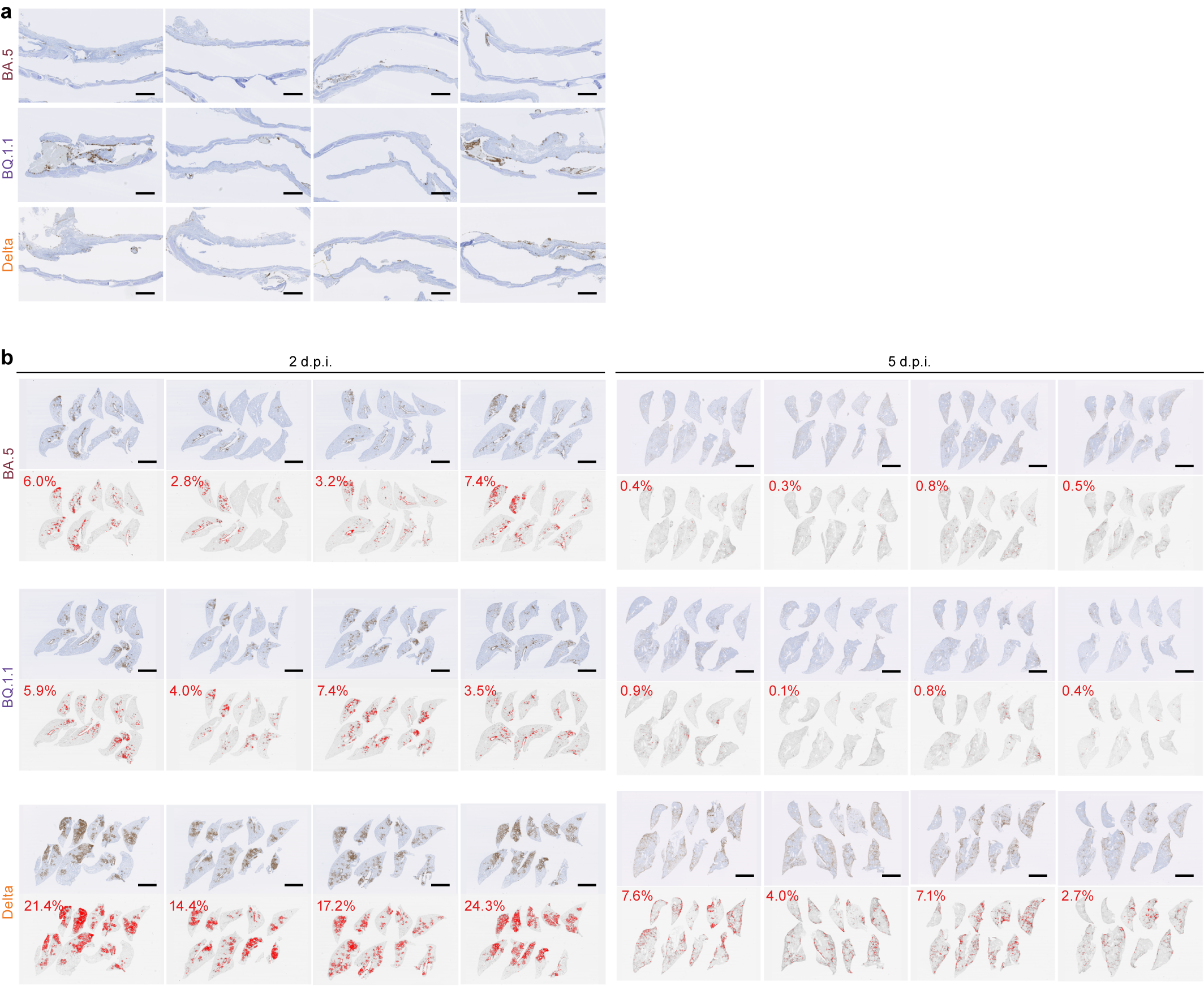
